# Carbonic anhydrase VII regulates dendritic spine morphology and density via actin filament bundling

**DOI:** 10.1101/736868

**Authors:** Enni Bertling, Peter Blaesse, Patricia Seja, Elena Kremneva, Gergana Gateva, Mari A. Virtanen, Milla Summanen, Inkeri Spoljaric, Michael Blaesse, Ville Paavilainen, Laszlo Vutskits, Kai Kaila, Pirta Hotulainen, Eva Ruusuvuori

## Abstract

Intracellular pH is a potent modulator of neuronal functions. By catalyzing (de)hydration of CO_2_, intracellular carbonic anhydrase (CA_i_) isoforms CAII and CAVII contribute to neuronal pH buffering and dynamics. The presence of two highly active isoforms suggests that they form spatially distinct CA_i_ pools enabling subcellular modulation of pH. Here we show that CAVII, unlike CAII, is localized to the filamentous actin network, and its overexpression induces formation of thick actin bundles and membrane protrusions in fibroblasts. In neurons, CAVII is enriched in dendritic spines, and its over-expression causes aberrant spine morphology. We identified amino acids unique to CAVII that are required for direct actin interactions, promoting actin filament bundling and spine targeting. Lack of CAVII in neocortical neurons leads to reduced spine density and increased proportion of small spines. Thus, our work demonstrates highly distinct subcellular expression patterns of CAII and CAVII, and a novel, structural role of CAVII.

## Introduction

Protons (H^+^ ions) have a strong modulatory effect on diverse cellular functions ranging from cell division (Roos & Boron 1981, Busa & Nuccitelli 1984) to directed cell movement (Patel & Barber, 2005; Van Duijn and Inouye, 1991). Within the central nervous system (CNS) neuronal excitability and signal transduction respond strongly to changes in intra- or extracellular pH (pH_i_ and pH_o_, respectively) due to the numerous, proton-sensitive molecular targets which include e.g. voltage- and ligand-gated ion channels (Traynelis and Cull-Candy, 1990; Pasternack et al., 1996; Tombaugh and Somjen, 1996; Duprat et al., 1997; Waldmann et al., 1997), gap junctions (Spray et al., 1981) and transmitter release (Sinning et al., 2011; Bocker et al., 2018). Although plasmalemmal acid-base transporters maintain neuronal pH_i_ close to 7.1 – 7.2 (Ruffin et al., 2014), deviations from this steady state level occur constantly. A particularly intriguing aspect in neuronal pH dynamics is that electrical activity can evoke intrinsic pH transients, which are either suppressed or enhanced by CA, depending on whether they are generated by transmembrane fluxes of H^+^ or HCO_3_^-^/CO_2_ fluxes, respectively (Taira et al., 1993; Voipio et al., 1995; Kaila and Chesler, 1998; Chesler, 2003; Sinning and Hubner, 2013).

The kinetics of neuronal pH_i_ changes depend both on the rate of plasmalemmal transporter function, and on the total intracellular buffering capacity β_t_ = β_i_ + β_CO2_. The intrinsic buffering power (β_i_) is mainly attributable to the titratable side chains of proteins. Hence, the presence of the highly mobile CO_2_/HCO_3_^−^ buffer system has, in addition to buffering (β_CO2_), an important role in enhancing diffusion of acid-base species within the cytoplasm (Voipio, 1998; Geers and Gros, 2000). The ability of the β_CO2_ system to operate rapidly is dictated by the presence of intracellular carbonic anhydrase activity (CA_i_) which accelerates the (de)hydration of CO_2_ to HCO_3_^−^ (Maren, 1967). The catalytically active cytosolic CA isoforms (SI, II, III, VII and XIII) are expressed in a cell-type specific manner. Neurons in the mature rodent hippocampus express two highly active CA_i_ isoforms, CAII and CAVII (Ruusuvuori et al., 2004; Ruusuvuori et al., 2013). In the central nervous system (CNS), CAVII localizes exclusively to neurons, and its expression starts at postnatal day (P) 10 - 12 in rodent hippocampal CA1 neurons (Ruusuvuori et al., 2004; Ruusuvuori et al., 2013). Notably, the catalytic product of the CO_2_/HCO_3_^−^ buffer system, HCO_3_^-^, acts as a major carrier of current in GABA_A_ receptor-mediated signaling (Kaila and Voipio, 1987), and the CAVII-driven replenishment of intraneuronal HCO_3_^-^ is a required for the development of paradoxical GABAergic excitation and network synchronization under conditions of prolonged activation of interneurons (Kaila et al., 1997; Ruusuvuori et al., 2004). After the expression of the housekeeping CAII isoform which commences in hippocampal pyramidal cells at around P20, both isoforms work in parallel to catalyze the intraneuronal CO_2_ (de)hydration (Ruusuvuori et al., 2013).

The co-expression of CAII and CAVII in neurons might point to distinct subcellular expression patterns of the two isoforms, based on specific interaction partners. While CA_i_s have not been reported to directly bind to cytoskeletal proteins, CAII has been reported to interact with membrane proteins. In cardiac ventricular myocytes, extramitochondrial CA_i_ activity-rich domains that facilitate CO_2_ movements (Schroeder et al., 2013) are generated by the interaction of CAII with the mitochondrial membrane and by the membrane-bound CAs localized to the sarcoplasmic reticulum (see also Wetzel et al., 2002; Scheibe et al., 2006). In erythrocytes, CAII interaction with the plasma membrane occurs possibly via binding to an integral membrane protein, the anion exchanger 1 (Vince and Reithmeier, 1998). Several other acid-base transporters have been proposed to act as structural and/or functional CAII interaction partners in expression studies (Sterling et al., 2001; Becker and Deitmer, 2007; Casey et al., 2009; Krishnan et al., 2015) (but see Boron, 2010). The formation of such CA complexes could conceivably allow generation of spatially distinct CA_i_ pools, which in neurons would gain further importance given their highly complex cellular architecture (Dong et al., 2015). Neurons face constant, spatially localized transmembrane fluxes of acid-base equivalents that are bound to cause local pH_i_ fluctuations, a detailed understanding of CA_i_ distribution is of great importance.

Here we show that, in contrast to CAII, CAVII is highly compartmentalized within neurons and that it has, in addition to its catalytic function, a role in F-actin dynamics. Specifically, we demonstrate that the distinct subcellular localization of CAVII is due to direct interactions with filamentous actin (F-actin). Further, we identify a novel role of CAVII in modulation of actin bundling that is not dependent on its catalytic activity. In neurons, CAVII is enriched in the actin-dense dendritic spines and CAVII depletion increases overall spine density, with a shift to smaller spine heads. These results show that neuronal CAVII has tightly linked functions in both cellular ion-homeostasis and cytoskeleton structure.

## Results

### CAVII binds to and bundles F-actin

To study the subcellular localization of CAII and CAVII we co-expressed EGFP-CAII and dsRed-CAVII fusion proteins in cultured NIH3T3 fibroblasts. The two isoforms showed strikingly different localization patterns. CAVII localized to cytosolic filamentous structures, whereas CAII was distributed homogenously throughout the cytoplasm and nucleus (Figure 1A). This difference prompted us to test whether CAVII co-localized with F-actin. Experiments, in which we expressed EGFP, EGFP-CAII or EGFP-CAVII in fibroblasts and stained F-actin with phalloidin-594 after fixation, showed no overlap of EGFP or EGFP-CAII with F-actin (Figure 1B-C). In contrast, EGFP-CAVII strongly co-localized with F-actin (Figure 1D). An exception to this were the outer edges of lamellipodia, in which EGFP-CAVII was not present (Figure 1-figure supplement 1). High levels of CA VII overexpression also caused marked changes in cellular morphology including the formation of thick, curving cytosolic stress fibers as well as filopodia-like plasmalemmal protrusions (Figure 1E). To compare the co-localization of the expressed CA proteins with F-actin in a quantitative manner, we measured the fluorescence intensities of EGFP and phalloidin-594 from cross sections through cells. These data, together with the correlation coefficients presented below in Figure 5, demonstrate that EGFP-CAVII, but not EGFP-CAII, co-localizes strongly with F-actin (Figure 1F-H).

**Figure 1.**
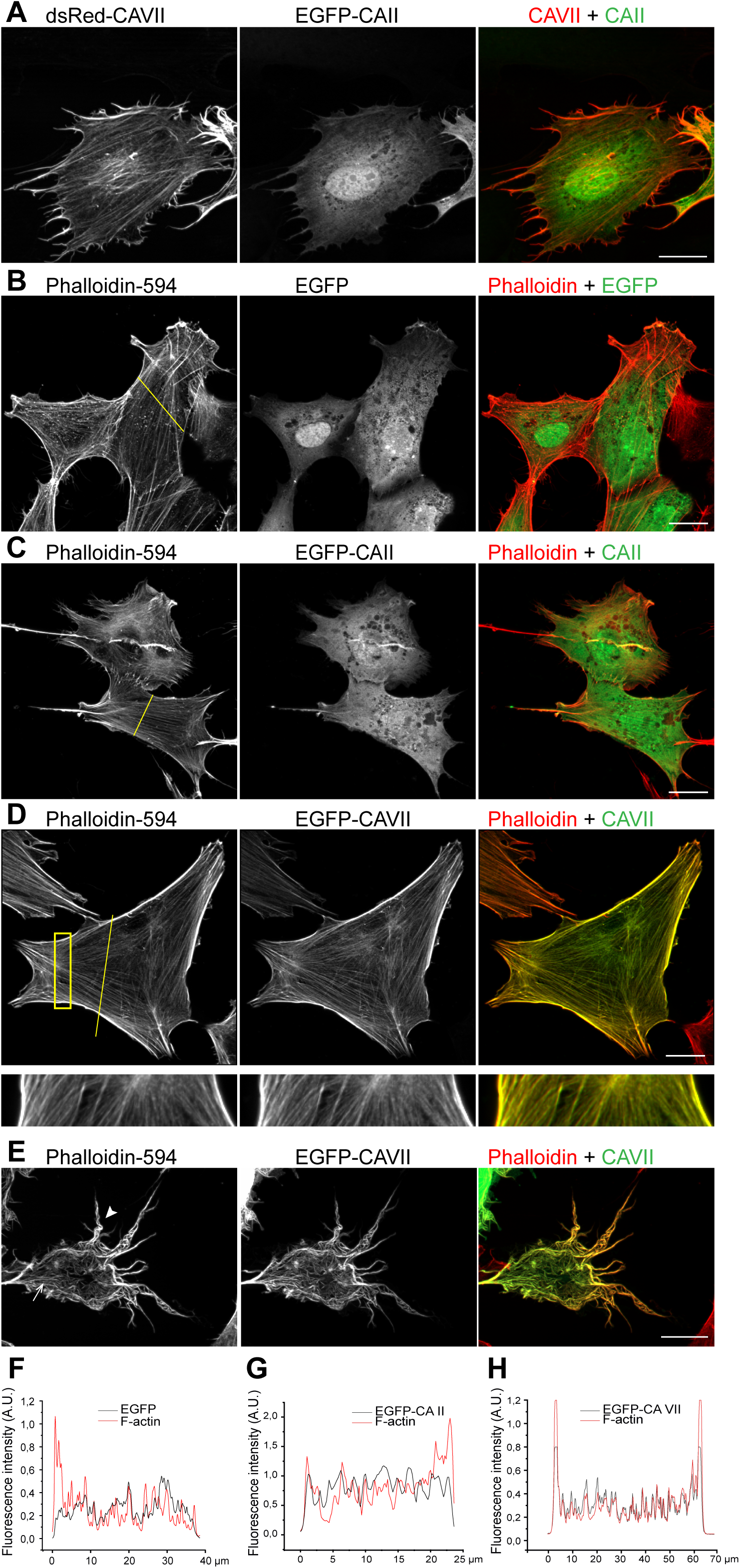
Subcellular localization of CAVII and CAII in fibroblasts. (A) NIH3T3 fibroblasts co-expressing dsRed-CAVII and EGFP-CAII (*n* = 4 independent replicates). Co-localization of EGFP and the two CA isoforms with filamentous actin studied in fibroblasts expressing (B) EGFP, (C) EGFP-CAII, or (D, E) EGFP-CAVII and stained with phalloidin-594 to visualize F-actin (*n* = 2, 7 and 9 independent replicates, respectively). Magnification of the area marked with the yellow rectangle in D shows co-localization of EGFP-CAVII with F-actin. (E) EGFP-CAVII caused a prominent overexpression phenotype with thick and curvy cytosolic actin bundles (arrow) and plasmalemmal protrusions (arrow head). (F-H) The normalized fluorescence emission intensity profiles for F-actin (red line) and (F) EGFP, (G) EGFP-CAII, or (H) EGFP-CAVII (black line). For the plots, pixel intensities were measured through the cross-section of the cell indicated by the yellow line in panels B-D. Scale bar in A-E 20 µm.

**Figure 5.**
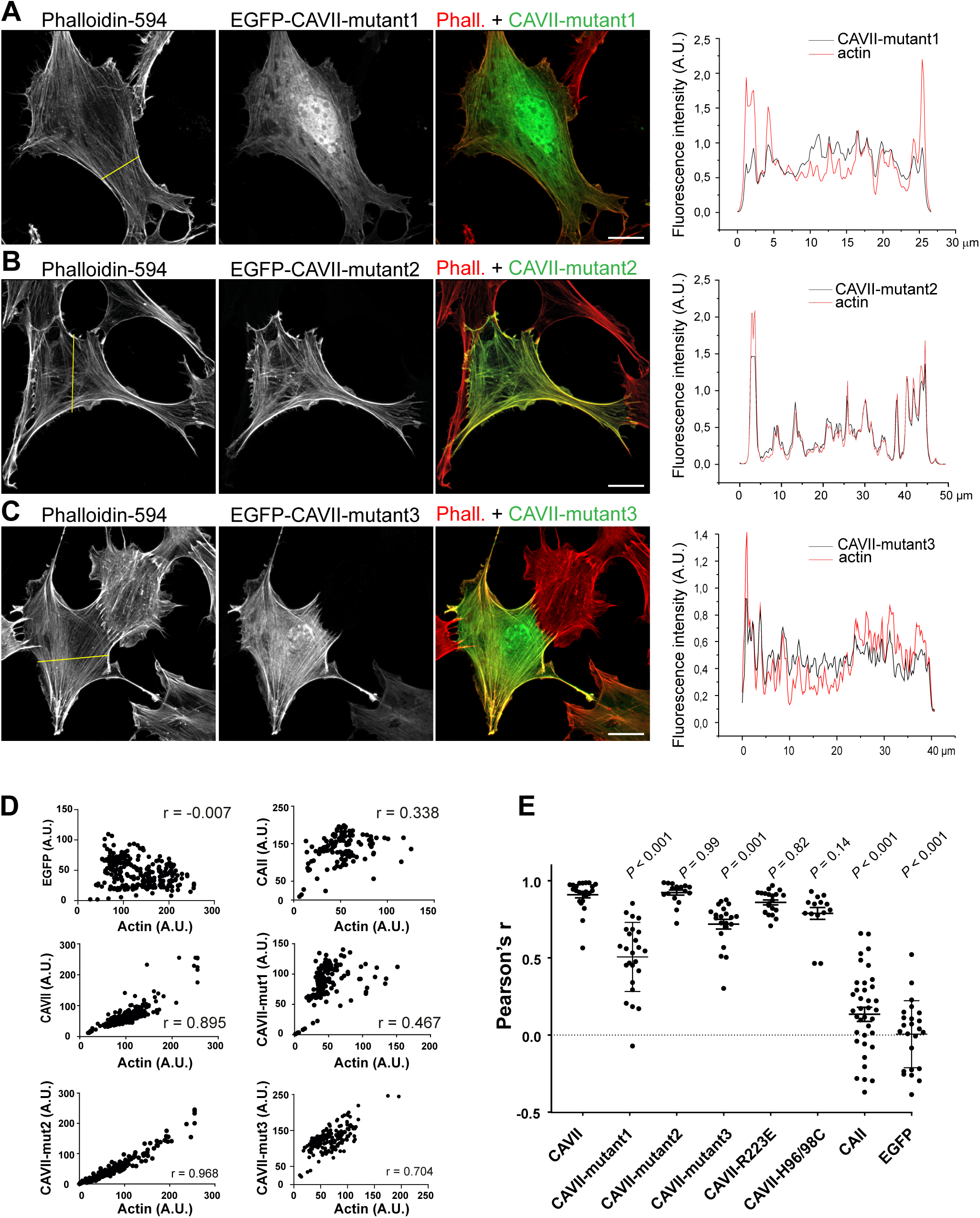
Subcellular localization of the chimeric CAVII constructs in fibroblasts. (A-C) NIH3T3 fibroblasts expressing EGFP-CAVII-mutant1 (A), EGFP-CAVII-mutant2 (B), and EGFP-CAVII-mutant3 (C). F-actin is visualized with Phalloidin-594. In the right-most panel of (A-C) are the normalized fluorescence intensity profiles of the mutated CAVII EGFP signal (black) and actin (red) and the yellow line in left-most panels indicates the cross-section from which the pixel intensities were measured. Scale bars 20 µm. (D) Analysis of the mutated CAVII and F-actin co-localization in cultured fibroblasts. Scatterplots of fluorescent intensities per pixel (EGFP vs. Phalloidin-594) along a cross section through a representative cell. Pearson’s correlation coefficient (r) for the analyzed cell is given in each panel. (E) Pearson’s correlation coefficient values calculated for the depicted constructs and compared to CAVII. F-actin had a strong positive correlation coefficient with EGFP-CAVII (r = 0.91 ± 0.02, *n* = 23 cells). Neither EGFP alone (r = 0.01 ± 0.04, *n* = 24) nor EGFP-CAII (r = 0.13 ± 0.05, *n* = 38) co-localized with F-actin (*P* < 0.001 for both constructs, when compared to CAVII). From the five mutated CAVII constructs, EGFP-CAVII-mutant1 (r= 0.56 ± 0.04, *P* < 0.001, *n* = 29) and EGFP-CAVII-mutant3 (r = 0.72 ± 0.03, *P* = 0.001, *n* = 21) co-localized less with F-actin actin when compared to CAVII. The co-localization of the other three mutated CAVII constructs, EGFP-CAVII-mutant2 (r = 0.92 ± 0.02, *P* = 0.9996, *n* = 18), EGFP-CAVII-R223E (r = 0.85 ± 0.02, *P* = 0.82, *n* = 21) and EGFP-CAVII-H96/98C (r = 0.79 ± 0.04, *P* = 0.14, *n* = 14) did not differ significantly from that of CAVII. Data are shown as mean ± SEM, statistical comparison against CAVII was done with one-way ANOVA and Dunnett’s multiple comparison test.

Next, we examined if the observed co-localization was based on a direct interaction between CAVII and F-actin. For these experiments, we used purified recombinant mouse CAVII protein (mCAVII), produced in the CHOEBNALT85 cell line. In an actin pull-down assay, mCAVII co-sedimented with F-actin, demonstrating a direct interaction between the two proteins (Figure 2A and Figure 2-figure supplement 1A). Interestingly, pH modulated this interaction: at pH 6.5, mCAVII bound F-actin with higher affinity than at pH 7.4 (Figure 2A and Figure 2-figure supplement 1A). CAII did not bind F-actin at either pH (Figure 2-figure supplement 1B,C).

**Figure 2.**
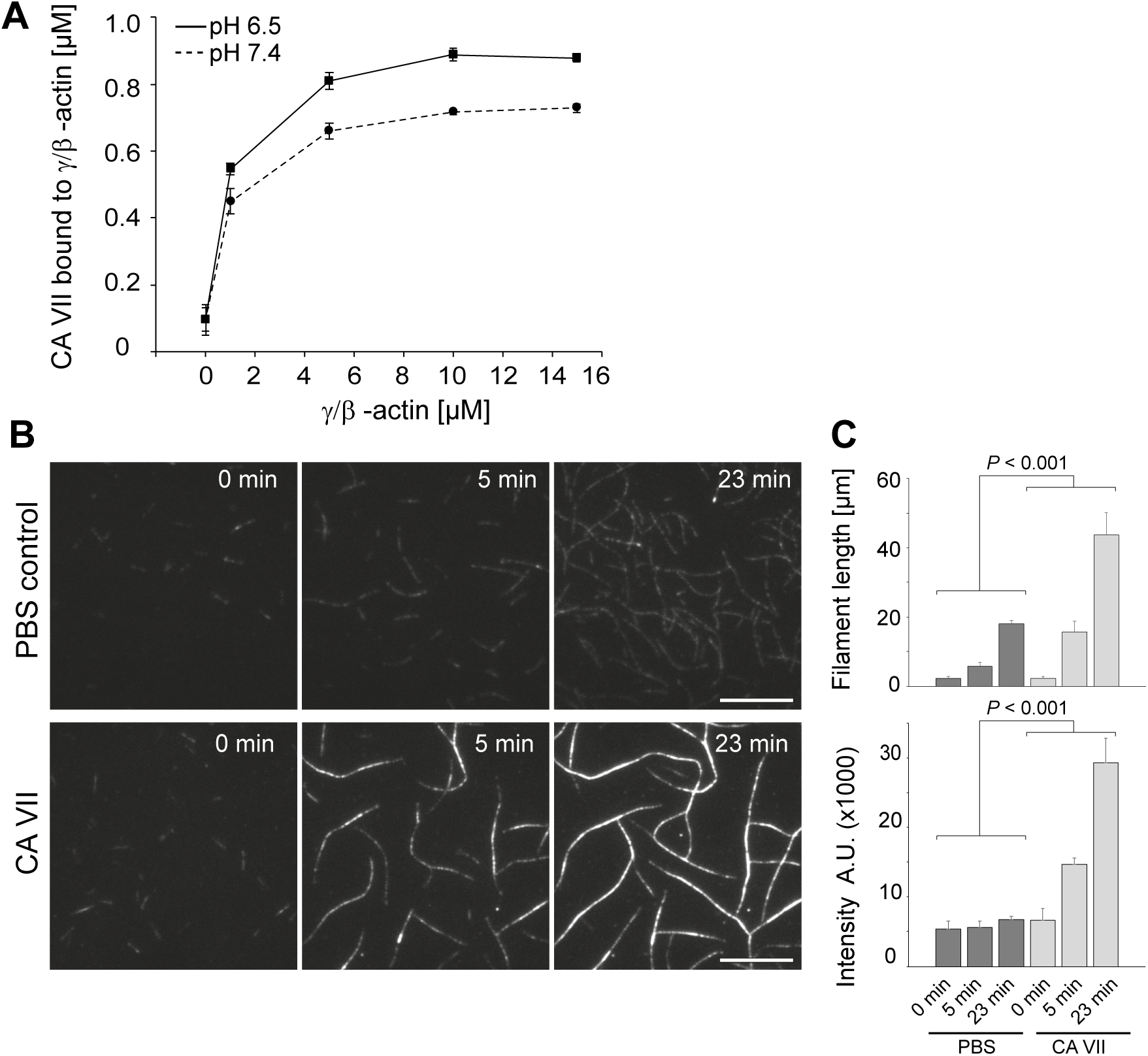
CAVII binds to filamentous actin and increases actin bundling. (A) Quantification of actin co-sedimentation assay shows that CAVII binds to F-actin. The binding is enhanced at more acidic pH (6.5 vs. 7.4) (*n* = 3 independent replicates at each actin concentration). (B) Fluorescence time-lapse images of F-actin bundling in an *in vitro* bundling assay. A mixture of unlabeled and Rhodamine labelled non-muscle actin was polymerized in the absence (PBS control, upper panel) or presence of mCAVII (lower panel). Numbers in images indicate the time after the onset of the experiment (0, 5 and 23 min). Scale bar 10 µm. (C) Quantification of the mean increase in filament length (*n* = 10 filaments at each time point) and relative fluorescence intensity (*n* = 30 - 31) in the absence and presence of mCAVII (1.12 µM). The data were analyzed using a general mixed model with time as a within unit factor and the presence of CAVII as a between unit factor. *n* = 3 independent repetitions, experiment repeats were included as a covariate and were non-significant. Data are presented as mean ± SD

As CAVII-expressing fibroblasts often showed abnormally thick actin structures, we set out to test if CAVII modulates actin bundling. To this end, we used an *in vitro* actin bundling assay in the absence and presence of 1.12 µM mCAVII (Figure 2B and Figure 2, Videos 1 and 2). Both actin filament length and fluorescence intensity of their cross section increased over time significantly in the presence of mCAVII when compared to the vehicle control (*P* < 0.001 for both length and intensity) (Figure 2C) indicating that CAVII had a profound effect on three-dimensional actin structures.

The ability of CAVII to increase bundling by crosslinking actin filaments implies a bivalent binding mechanism that conventional actin cross-linkers, such as α-actinin and fimbrin, most often achieve through homodimerization (Puius et al., 1998). We analyzed the oligomeric status of mCAVII expressed as a secreted construct with analytical size exclusion chromatography and multi-angle light scattering (SEC-MALLS) (Figure 2-figure supplement 2). Majority of the protein is monomeric at the tested 33 µM concentration (estimated molecular mass of approximately 35-42 kDa), matching quite well with the theoretical size of monomeric CAVII (30.5 kDa). A molecular weight of approximately 53-75 kDa was determined for a minor peak eluting at an earlier time point, which corresponds to a putative CAVII dimer.

Based on the above results, we tested if CAVII-mediated actin bundling stabilizes actin filaments. This was done by exposing transfected cells to latrunculin B which sequesters free actin monomers and can thus be used as a robust method to quantify the rate of actin filament depolymerization (Figure 3). The experiments showed that in control cells, expressing dsRed, actin filaments depolymerized within a few minutes (Figure 3A). In contrast, over 60 % of the dsRed-CAVII expressing cells had visible F-actin structures even after 30 minutes of latrunculin B treatment (Figure 3B).

**Figure 3.**
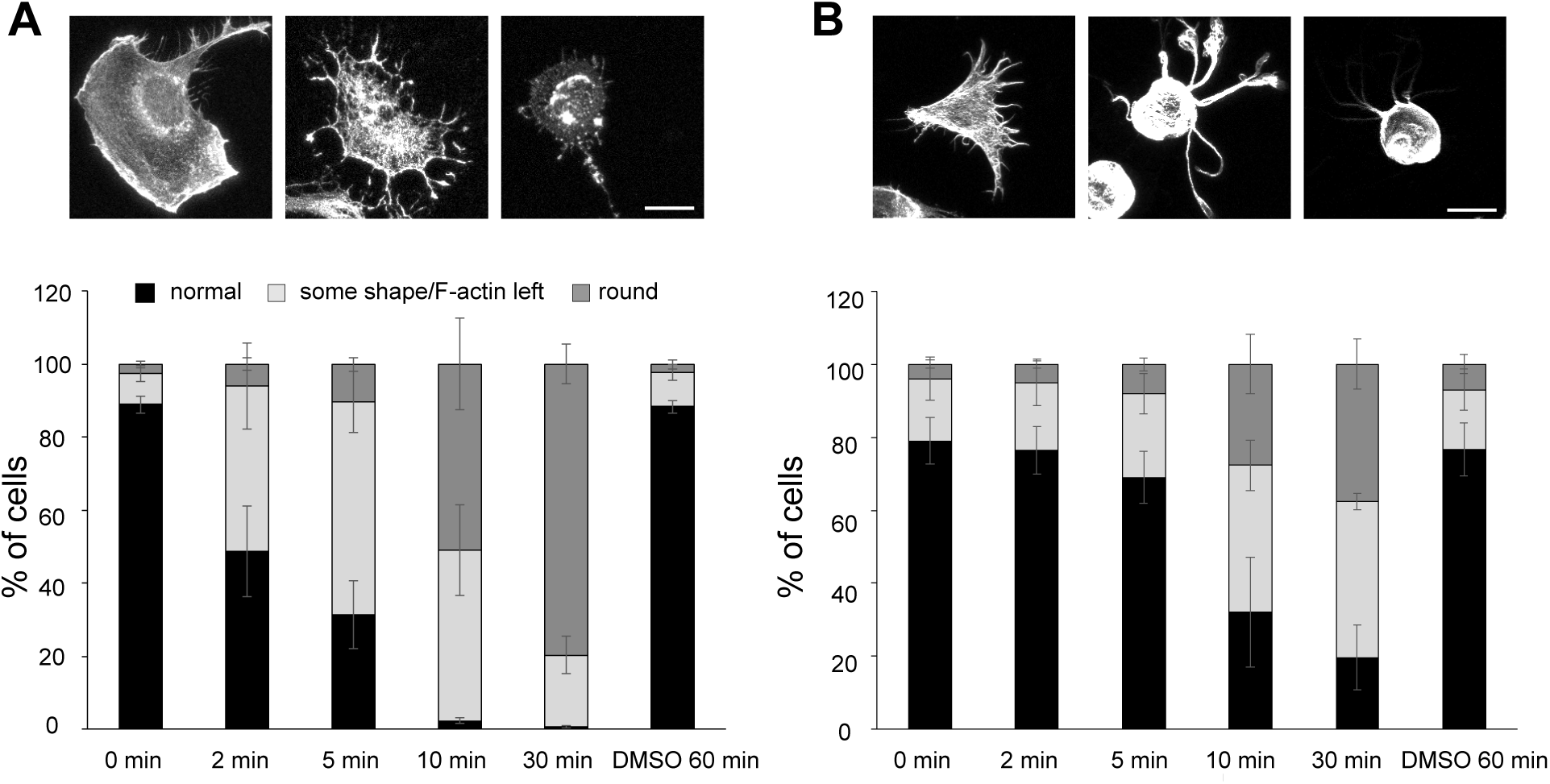
CAVII over-expressing NIH3T3 are resistant to the latrunculin B treatment. NIH3T3 cells transfected with dsRed (A) or dsRed-CAVII (B) were incubated in growth medium with 5 µM Latrunculin B for 0, 2, 5, 10, or 30 minutes, or in an equal amount of DMSO for 60 minutes. Analyses of experiments show that in cells transfected with dsRed-CAVII F-actin structures collapse more slowly than in the dsRed-transfected ones. For the analysis, cells were categorized to three groups as “normal”, “some shape/F-actin left” and “round”. The higher panel shows example images of the cells in all three categories for (A) dsRed- or (B) dsRed-CAVII transfected cells (actin visualized with Phalloidin-488). A hundred cells per each time point from each experiment were counted and categorized. Scale bar 50 µm

These data demonstrate that CAVII directly binds to and bundles F-actin, and that CAVII over-expression stabilizes existing actin filaments.

### The CAVII surface motif DDERIH is crucial for actin binding

As no previously known actin-binding domains (Paavilainen et al., 2004; Lee and Dominguez, 2010) are present in CAVII, we sought to identify which CAVII sequence features contribute to actin binding. We first analyzed publicly available CAII and CAVII protein structures (Eriksson et al., 1988; Di Fiore et al., 2010) to identify potential actin-interaction surfaces. These comparisons revealed a structural motif at the CAVII protein surface that clearly differs from CAII (Figure 4). The amino acids 101-105, 113, 115 and 237-242 are widely distributed in the primary sequence of CAVII, but form a continuous “ridge” at the protein surface (Figure 4A,B). Superimposing the CAVII α-helix 6 on the twinfilin-C/G-actin structure (Paavilainen et al., 2007) suggested that CAVII α-helix 6 including the amino acids 237-242 could also be a putative interaction site (Figure 4C). Arginine (R223) located to α-helix 6 has a positive charge that is reversed in CAII. Thus, we hypothesized that this amino acid might have a role in CAVII/actin interaction. The alignment of the protein sequences of CAII and CAVII confirmed that these amino acids are not conserved between the two isoforms (Figure 4D). In addition, a sequence comparison with the human cytosolic CA isoforms revealed that these motifs are unique to CAVII (Figure 4-figure supplement 1).

**Figure 4.**
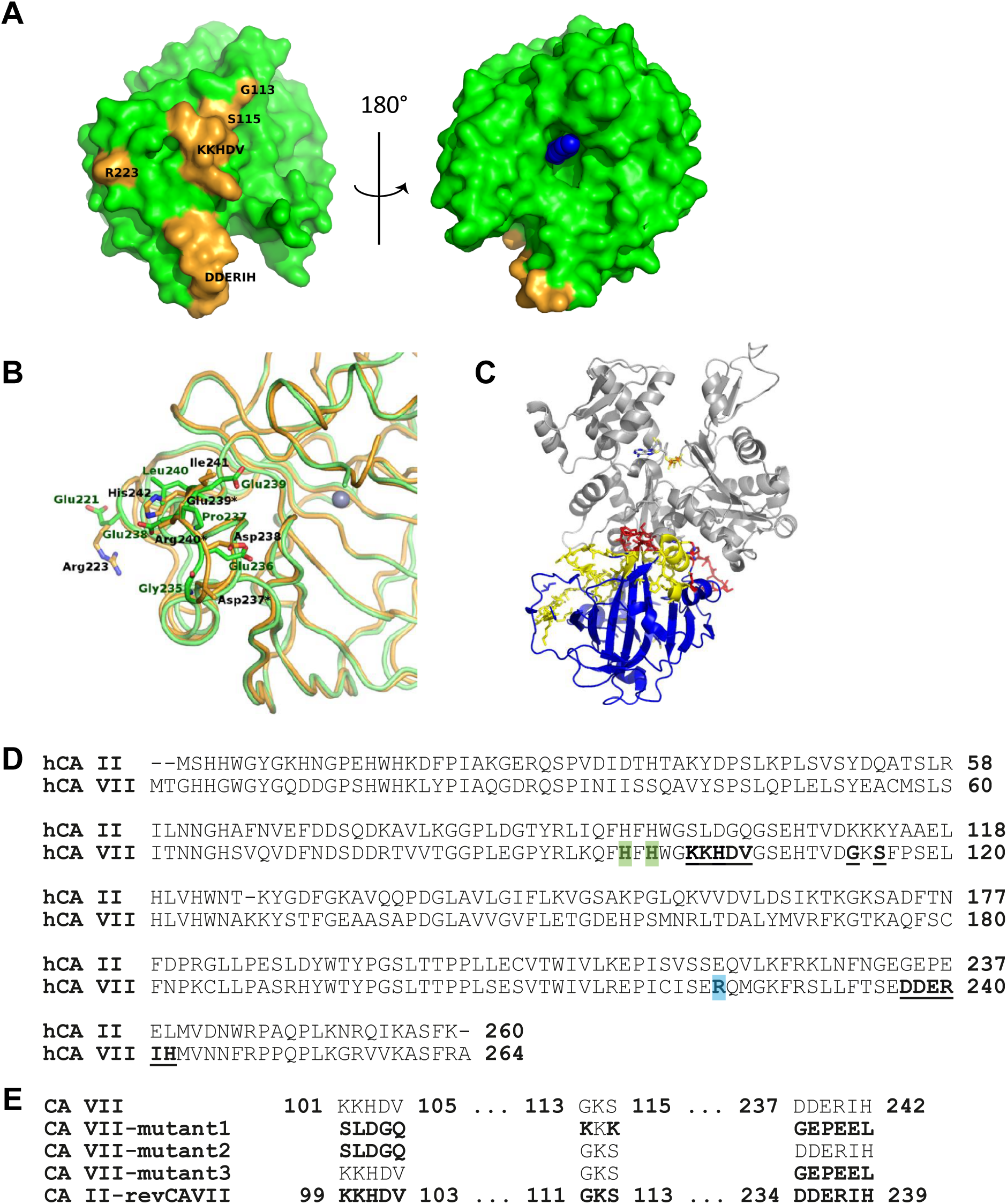
Structure and sequence comparison of CAVII and CAII. (A) Three-dimensional representation of the CAVII structure. The amino acids 101-105, 113, 115 and 237-242 form a ridge at the protein surface (highlighted in yellow). R223 is located close the ridge. The blue spherical structure in the 180° view (right panel) depicts the inhibitor 6-ethoxy-1,3-benzothiazole-2-sulfonamide bound to the active site. (B) Overlay of CAII and CAVII three-dimensional structure illustrating structural differences at the protein surface. Note the prominent position of R223 in CAVII. The main chain, side chains of selected residues, and zinc ion are shown as ribbon, sticks, and sphere representation, respectively. Carbon atoms of CAII and CAVII are colored green and orange, respectively. The side chains of Asp237, Glu239, and Arg240 (labeled with *) of CAVII are not defined by the electron density and thus omitted from the published PDB file (3MDZ). (C) Superimposing the CAVII α-helix-6 (CAVII in blue, areas where mutations1-3 are located in yellow, and the putative actin interactin CAVII helix in red) on the Twf-C/G- actin structure (grey) shows a sterically compatible structure. (D) Sequence alignment of human CAVII and CAII protein sequences generated using the Clustal O (1.2.1) multiple sequence alignment. Residues forming a ridge at the CAVII protein surface in the CAVII 3D structure are highlighted in the CAVII sequence (bold/underlined). R223 is marked in turquoise and H96 and H98 (mutated to gain a catalytically loss-of-function mutant) are highlighted in green. (E) Schematic representation of four mutants with a full (mutant1) or partial (mutant2 and mutanteplacement of the amino acids encoding the ridge in CAVII by the corresponding CAII sequence. In the reversed mutant (CAII-revCAVII) the amino acids replaced in mutant1 were introduced to CAII. Panels A and B were prepared using PyMOL (PyMOL The PyMOL Molecular Graphics System, Version 1.4.1 Schrödinger, LLC.).

To explore the putative CAVII-actin binding domains, we generated a set of CAVII mutants in which subsets of the amino acids mentioned above were replaced with the corresponding amino acid sequences of CAII (Figure 4E). The mutants were expressed in NIH3T3 cells as EGFP fusion proteins and their subcellular distribution and co-localization with F-actin were examined after staining cells with phalloidin-594 (Figure 5A-C and Figure 5-figure supplement 1). When amino acids forming the “ridge” (101-105, 113, 115 and 237-242) were replaced with the corresponding amino acids of CAII (EGFP-CAVII-mutant1; see Figure 4D,E), the strict co-localization with F-actin was abolished and the construct did not induce detectable changes in the actin structure or cell morphology (Figure 5A). To characterize the interaction motif in more detail, we separately mutated the amino acids 101-105 (KKHDV; EGFP-CAVII-mutant2) or 237-242 (DDERIH; EGFP-CAVII-mutant3). The localization of EGFP-CAVII-mutant2, with only the KKHDV motif replaced by the corresponding CAII sequence, was unchanged, when compared to EGFP-CAVII. EGFP-CAVII-mutant2 co-localized strongly with F-actin (Figure 5B), and the over-expression phenotype with thick, curving cytosolic actin bundles and plasmalemmal protrusions (Figure 5-figure supplement 1A) was similar to that seen with EGFP-CAVII. Interestingly, replacement of the DDERIH motif alone (EGFP-CAVII-mutant3) reduced the co-localization with F-actin, and expression of the mutant did not visibly affect cellular actin structures (Figure 5C). We compared the co-localization of the different EGFP-tagged variants with phalloidin-594 by plotting the fluorescence intensities per pixel against each other (shown in right-most panels in Figures 5A-C and Figure 5-figure supplement 1E-G) and calculating Pearson’s correlation coefficient (Figure 5D,E). Comparison of the scatter plots from individual cells revealed that, firstly, F-actin had a strong positive correlation coefficient with CAVII, but not with CAII. Secondly, the correlation coefficients of EGFP-CAVII-mutant1 and EGFP-CAVII-mutant3, but not that of EGFP-CAVII-mutant2, were reduced when compared to CAVII (Figure 5E). This indicates that the DDERIH motif forms an important part of the CAVII/actin interaction site.

Despite the changes seen with EGFP-CAVII-mutant1 and EGFP-CAVII-mutant3, an analogue mutant of CAII, containing all mutations of EGFP-CAVII-mutant1 in a reversed manner (EGFP-CAII-revCAVII, see Figure 4E), maintained the diffuse cytosolic localization pattern seen with CAII (Figure 5-figure supplement 1B,E).

When the positively charged arginine 223, which we hypothesized to play a role in CAVII-actin binding, was mutated to a negatively charged glutamic acid (EGFP-CAVII-R223E), co-localization with F-actin was not affected (Figure 5E and Figure 5-figure supplement 1C,F), but the generation of thick cytosolic actin bundles and plasmalemmal protrusions was reduced. Finally, to examine whether CAVII catalytic activity affects actin co-localization, we expressed a functionally inactive mutant, EGFP-CAVII-H96/98C (Kiefer and Fierke, 1994), in NIH3T3 cells. This mutant retained its co-localization with actin (Figure 5E and Figure 5-figure supplement 1D,G), suggesting that enzymatic activity is dispensable for the interaction with actin. The different impact of the various constructs on actin cytoskeleton morphology did not seem to correlate with overall construct expression levels (Figure 5-figure supplement 2). EGFP-CAVII-R223E, for example, had comparable actin co-localization and expression levels to EGFP-CAVII, but did not induce thick actin bundles.

Together, these comparisons provide compelling evidence for structural motifs consisting of widely spread amino acids within the primary structure of CAVII and containing the DDERIH sequence as the CAVII/actin binding site.

### CAVII localizes to dendritic spines and its overexpression disrupts spine morphology

We next moved on to study the subcellular localization of CAII and CAVII in rat hippocampal neurons by co-expressing dsRed-CAVII and EGFP-CAII in neuronal cultures. While EGFP-CAII homogenously distributed along the somato-dendritic axis, dsRed-CAVII was prominent in the actin-rich spines (Figure 6A). Co-expression of mCherry-actin and EGFP-CAII further confirmed that CAII, similar to EGFP alone, localizes diffusely to dendrites and spines and causes no apparent changes in the structure of dendritic spines (Figure 6-figure supplement 1A,B). In contrast, the EGFP-CAVII distribution overlapped with that of mCherry-actin and, in line with the CAVII-dependent modulation of actin structures detected in fibroblasts, it had a marked effect on spine morphology: CAVII-expressing neurons had a high proportion of aberrant spines, i.e. thick, filopodia-like dendritic protrusions with no clear spine head (Figure 6-figure supplement 1C). The effects of EGFP, EGFP-CAII and EGFP-CAVII on spine density and structure in cultured neurons are summarized in Figure 6-figure supplement 1I.

**Figure 6.**
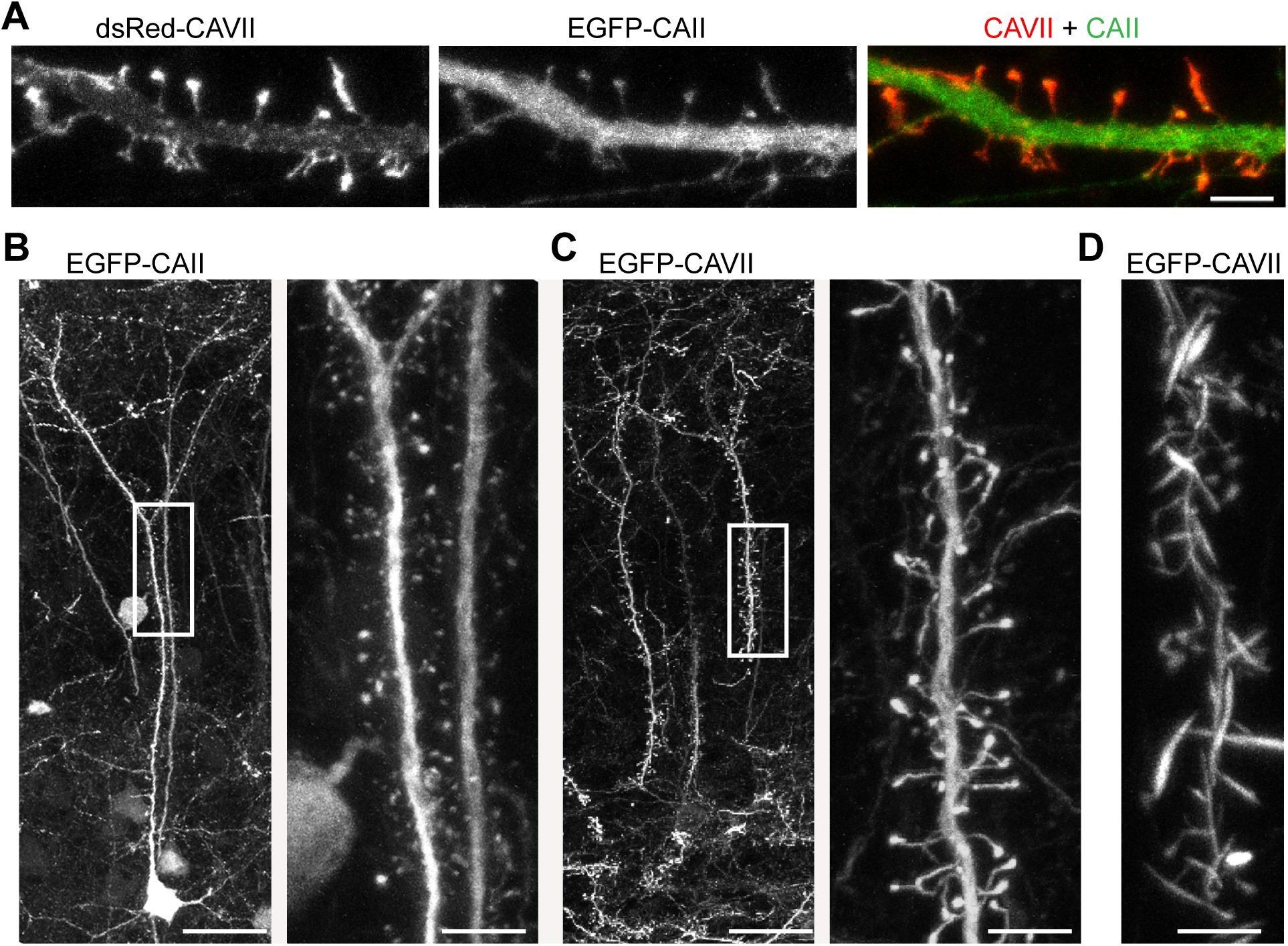
Localization of CAII and CAVII in neurons. (A) Isoform-specific subcellular localization shown in cultured hippocampal neurons (DIV14) co-expressing dsRed-CAVII (*left*) and EGFP-CAII (*middle*). Representative confocal images of precocious *in vivo* expression of (B) EGFP-CAII and (C) EGFP-CAVII in P40 mouse cortical layer 2/3 pyramidal neurons. Neurons were transfected at E14.5 with EGFP-CAII or EGFP-CAVII using *in utero* electroporation and images were taken from fixed slices. Right panels in B and C show higher magnification of the primary apical dendrite marked with a box. (D) The expression of EGFP-CAVII disrupted the normal spine morphology and induced the formation of thick, filopodia-like protrusions. *n* = 5 independent repeats for cultured neurons and two animals/construct *in vivo*. Scale bar in (A) 5 µm; (B and C): 5 µm, insets in B, C and panel D: 25 µm

Of the mutant proteins, EGFP-CAVII-mutant2 co-localized with mCherry-actin and caused a similar change in spine morphology as CAVII (Figure 6-figure supplement 1D). When the DDERIH motif was mutated individually (EGFP-CAVII-mutant3), or together with the other amino acids forming the ridge (EGFP-CAVII-mutant1), the subcellular distribution of the fusion proteins was homogenous along dendrites and dendritic spines, and expression of these constructs had no obvious effect on the spine morphology (Figure 6-figure supplement 1E,F). EGFP-CAVII-R223E and EGFP-CAVII-H96/98C, both of which co-localized with F-actin in fibroblasts, showed strongly overlapping localization with mCherry-actin in dendritic spines (Figure 6-figure supplement 1G,H).

Finally, to study the localization of CAVII and CAII in neurons *in vivo,* we expressed EGFP-CAVII or EGFP-CAII in cortical layer 2/3 pyramidal neurons using *in utero* electroporation and examined transfected neurons in fixed slices from P40 mice. We saw that, compared to EGFP-CAII (Figure 6B), EGFP-CAVII localized strongly to dendritic spines and induced the formation of abnormal, filopodia-like dendritic protrusions (Figure 6C,D) which is in line with the observations made in cultured neurons.

### Genetic deletion of CAVII changes cortical layer 2/3 pyramidal neuron spine density and morphology *in vivo*

The spine-targeted expression of CAVII and its interaction with the actin cytoskeleton raised the question whether genetic deletion has an effect on dendritic spines. For this, we did electrophysiological recordings and structural analysis of neurons using CAVII knockout (CAVII KO) and wild type (WT) mice (Ruusuvuori et al., 2013). Since spines are the major site for excitatory synaptic input, we measured miniature excitatory postsynaptic currents (mEPSCs) from WT and CAVII KO somatosensory cortex layer 2/3 pyramidal neurons (Figure 7A). Neither mEPSC amplitude (14.93 ± 1.16 pA vs. 14.71 ± 1.10 pA, *P* = 0.90) nor frequency (5.37 ± 1.62 Hz vs. 6.70 ± 2.24 Hz, *P* = 0.63) differed between the genotypes (*n* = 7 WT and *n* = 4 CAVII KO neurons from five WT and four CAVII KO mice). Structural analysis was done from Lucifer Yellow (LY) labeled somatosensory cortex layer 2/3 pyramidal neurons from P34 - P37 WT and CAVII KO mice (Figure 7B) as described earlier (Fiumelli et al., 2013). Interestingly, genetic ablation of CAVII significantly changed dendritic architecture *in vivo* (Figure 7C). In CAVII KO neurons (8279 spines analyzed from 28 neurons/four animals) spine density on the second order apical and basal dendritic shafts increased by 38 ± 16 % and by 42 ± 16 % (*P* < 0.001), respectively, in comparison to the WT neurons (8730 spines from 30 neurons/two animals). The change in spine density was due to a specific increase in immature-type of spines with small spine heads. The average spine head diameter in CAVII KO neurons was 0.37 ± 0.01 μm (421 spines from 15 neurons) and in WT neurons 0.48 ± 0.01 μm (467 spines from 15 neurons). Compared to WT, the distribution was shifted significantly towards smaller spine heads in the CAVII KO neurons (W = 134540, *P* < 0,001, Wilcoxon rank sum test) (Figure 7D).

**Figure 7.**
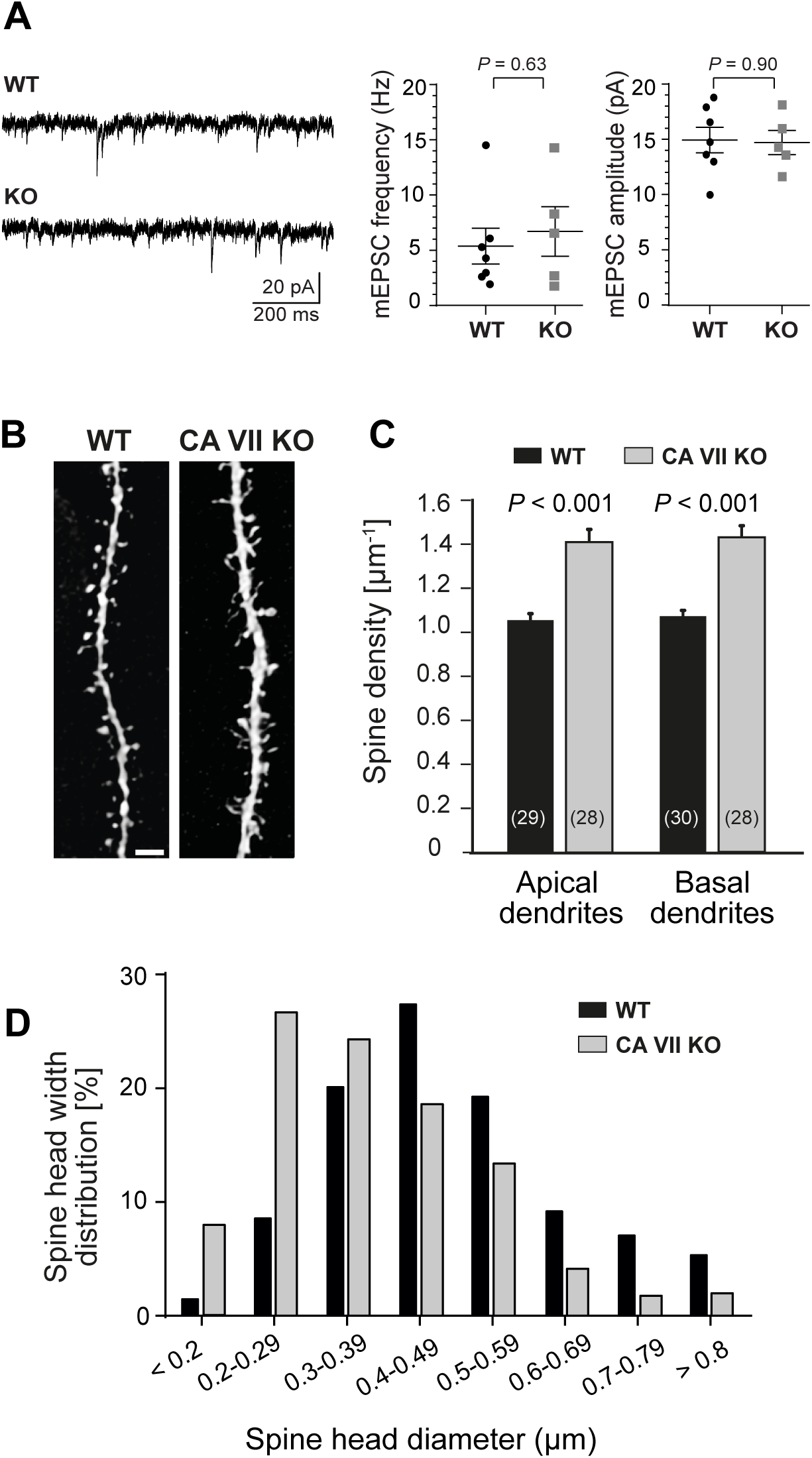
Layer 2/3 cortical pyramidal neurons in CAVII KO mice have high dendritic spine density and smaller spines but mEPSC frequency and amplitude are not affected. (A) Comparison of mEPSCs in cortical layer 2/3 pyramidal neurons from P30 – P40 WT and CAVII KO mice. Sample traces of mEPSC recordings from WT and CAVII KO neurons, low-pas filtered at 1 kHz (left). The data are summarized in the bar diagrams (right). mEPSC frequency (*P* = 0.63) and amplitude (*P* = 0.90) were not significantly different between CAVII KO and WT neurons (*n* = 4 and 7 neurons, respectively, Student’s independent samples *t*-test). (B) Representative confocal images of apical dendrites from Lucifer Yellow injected cortical layer 2/3 pyramidal neurons from WT and CAVII KO mice. The dendritic spine density and spine head size were examined in fixed slice preparations from P34 - P37 mice. Scale bar 2 µm. (C) Summary of the spine density analysis done from the Lucifer Yellow injected neurons. A total of 8279 spines were analyzed from the CAVII KO mice (*n* = 4 animals), and 8730 spines from the WT control mice (*n* = 2 animals). Number of analyzed cells is indicated in brackets in the bar diagram. Mann-Whitney test and Student’s *t*-test with Welch-correction were used for statistical analysis for apical and basal dendrites, respectively. (D) The spine head width distribution, measured from 467 spines from 15 WT cells and 421 spines from 15 CAVII KO cells, differed significantly between the genotypes (Wilcoxon rank sum test with continuity correction, W = 134540, *P* < 0.001). Data in B and C are given as mean + SEM.

## Discussion

The two cytosolic CA isoforms present in mature rodent central neurons are both catalytically highly active (Earnhardt et al., 1998) and have a high sequence and structural similarity (Di Fiore et al., 2010). Here we show that, despite these similarities, CAVII has unique characteristics, profiling it as a novel, multifunctional protein within the CNS. We demonstrate a prominent difference in the subcellular localization of the isoforms, which is based on a pH-dependent interaction of CAVII and actin. In neurons, CAVII is enriched in the actin-dense dendritic spines and affects the actin cytoskeleton thus altering both dendritic spine morphology and density.

### The subcellular distribution of CAVII is dictated by its interaction with F-actin

In cultured fibroblasts, expression of dsRed- and EGFP-fusion constructs of CAVII and CAII revealed a mutually exclusive subcellular localization. CAVII is present in the immediate vicinity of actin filaments, in contrast to the diffuse cytoplasmic localization of CAII. In hippocampal neuronal cultures, as well as in cortical neurons transfected *in vivo,* CAVII shows a preferential spine-targeted localization, whereas CAII distributed evenly in the cytosol over the somato-dendritic axis. The diffuse cytoplasmic distribution of CAII is well in line with previous localization results from non-erythroid cells (Wang et al., 2002; Stridh et al., 2012; Al-Samir et al., 2013) and fits with the idea that this ubiquitous and high-activity isoform serves a housekeeping role in cytosolic pH buffering. Homogenously distributed, soluble CA can efficiently dissipate cytosolic acid-base gradients (Voipio, 1998; Stewart et al., 1999; Boron, 2010). Interestingly, a recent study on cardiac myocytes shows that the majority of nuclear pH buffering is sourced from the cytoplasm in the form of mobile buffers (Hulikova and Swietach, 2015), motivating further work on the localization of CAII and its possible role in pH-dependent regulation of transcription (Bumke et al., 2003; Neri and Supuran, 2011).

Our main finding is that the subcellular distribution of CAVII is dictated by its binding to actin, in particular to F-actin, which is a novel property with regard to all cytosolic CAs studied to date. Furthermore, CAVII interacts only with a specific subset of actin filaments. EGFP-CAVII strongly co-localized with F-actin in fibroblast stress-fibers and neuronal dendritic spines, but the edges of the highly dynamic fibroblast lamellipodia, consisting mainly of branched actin, were largely devoid of CAVII. Given the key role of CAs in the modulation of pH, a particularly interesting finding is that the CAVII–actin interaction is pH-sensitive and enhanced at acidic pH (6.5 vs. 7.4). Compared with the actions of the previously known pH-sensitive actin binding proteins (see below), CAVII could thus counteract gelsolin-dependent severing of F-actin, which takes place upon acidification (Lagarrigue et al., 2003). Together with our latrunculin data these results indicate that CAVII could stabilize actin structures when the intracellular compartment is acidified, which occurs under various pathophysiological conditions such as stroke (Pavlov et al., 2013) and epilepsy (Siesjö et al., 1993). Furthermore, an increase in neuronal activity subjects brain cells to surges of lactate produced by the glycolytic pathway (see Yellen, 2018). Within neurons, CAVII is ideally located to facilitate H^+^-coupled lactate (Halestrap, 2013) efflux across the neuronal plasma membrane (Yellen, 2018).

CAVII not only binds to F-actin but it also increases actin bundling *in vitro*. Our bundling assay was done in the nominal absence of CO_2_/HCO_3_^-^ indicating that the CAVII-dependent enhancement of bundling is not dependent on the catalytic activity of CAVII. This is well in line with our result that the enzymatically inactive H96/98C CAVII mutant maintains it co-localization with F-actin. Notably, the enzymatic activity of CAVII is much more sensitive to the concentration of chloride than that of CAII (Vullo et al., 2006). It is therefore possible that in highly compartmentalized structures subjected to large ionic fluctuations, such as dendritic spines (Rose et al., 1999; Brini et al., 2017), CAVII’s catalytic activity is compromised while its interaction with actin is maintained. The presence of neuronal CAII in spines would thus contribute to CO_2_/HCO_3_^-^ based buffering even when the ionic milieu is transiently changed.

Our time-lapse imaging experiments visualizes the CAVII-dependent change in actin dynamics (SI Videos 1 and 2). The rapid assembly/disassembly of actin filaments seen under control conditions was suppressed in the presence of CAVII and the generated actin-bundles were thicker and more stable. Based on the SEC-MALLS results, CAVII-dependent bundling may be achieved, at least partly, through CAVII homodimerization. In the intracellular milieu, even a small proportion of free dimers might be enough if actin acts as a sink for the CAVII dimers thereby shifting the dimerization process to the right.

### CAVII-actin interaction induces morphological changes

The changes detected in the actin cytoskeleton of CAVII-expressing cells are consistent with the biochemical assay data and show that CAVII modulates higher-order actin structures in the cytoplasm. Fibroblasts transfected with CAVII generate numerous filopodia-like protrusions projecting from the cell surface, and have thick, sometimes curving, cytosolic stress fibers. In neurons overexpressing CAVII, both in cell cultures and *in vivo,* dendritic spines eventually lose their morphological diversity (categorized as thin, stubby, and mushroom spines according to (Bourne and Harris, 2008; Hotulainen and Hoogenraad, 2010)) and turn into thick protrusions, which lack a clear spine head. Similar filopodia structures sprout even from the neuronal cell body. These phenotypic characteristics closely resemble those seen with the brain-specific actin-bundling protein drebrin-A in fibroblasts (Shirao et al., 1994) and in cultured neurons (Hayashi and Shirao, 1999; Mizui et al., 2005). The disturbed spine morphology shows that the actin network normally forming these structures is modified to more rigid actin bundles. Together with the biochemical bundling assay results, the CAVII overexpression phenotype suggests that CAVII has a stabilizing effect on F-actin. We tested this hypothesis in experiments where fibroblasts were exposed to the actin polymerization inhibitor latrunculin B. Compared to control cells, fibroblasts expressing CAVII maintained F-actin structures significantly longer, confirming a direct stabilizing effect of CAVII.

### Identification of the CAVII – actin interaction site

When Montgomery et al. (1991) first identified the human CAVII, they recognized several poorly conserved regions that were predicted to “*be located towards the surface of the protein”.* Our work shows that one of these regions, residues 232 – 248 encoded by exon 7, is critically involved in CAVII-actin interaction. When we replaced amino acids 237 – 242 (DDERIH) with corresponding amino acids from CAII either alone (EGFP-CAVII-mutant3) or together with additional mutations (amino acids 101-105, 113 and 115; EGFP-CAVII-mutant1), co-localization of F-actin and the mutated EGFP-CAVII proteins decreased compared to EGFP-CAVII. Replacing the corresponding sequence of CAII by the CAVII amino acids forming the ridge (101-105, 113, 115 and 237-242) was not sufficient to induce an actin-binding phenotype of CAII, pointing to a complex three-dimensional structure of the actin-binding site. The catalytically inactive CAVII-H96/98C mutant and CAVII with the point mutation R223E, disrupting the positive charge at the CAV II α-helix-6 (αG), still bound actin, but formation of thick cytosolic actin bundles was suppressed.

### Genetic deletion of CAVII changes spine density and morphology

Since ectopic expression of CAVII had such a prominent effect on spine morphology, we examined how deletion of CAVII affects dendritic spines *in vivo*. The increase in spine density, together with the shift to smaller spine head size in CAVII KO mouse cortical neurons, demonstrates that endogenous CAVII serves a structural role in dendritic spines. As our quantification in these experiments is based solely on the head diameter (Bourne and Harris, 2008), we cannot address how the proportion of filopodia and thin spines changes. However, it is well established that spines with small spine heads i) are short-lived and dynamic structures (Holtmaat et al., 2005); ii) are more abundant in early development and iii) make only occasional contacts with presynaptic terminals (Berry and Nedivi, 2017). It is thus not surprising that the detected increase in small spines (spine heads < 0.3 µm) in CAVII KO cortical neurons did not alter the basic pre- or postsynaptic properties, measured as the frequency and amplitude of mEPSCs, respectively. It is tempting to suggest that the overproduction of small spines, actively searching for presynaptic partners, renders CAVII KO neurons in “a more juvenile” morphological state. Previous *in vivo* studies have demonstrated that there are pools of spines with different turn-over rates and these pools change over the postnatal development and differ greatly between brain areas even in mature animals (Holtmaat et al., 2005; Attardo et al., 2015; Pfeiffer et al., 2018). Hence, it would be intriguing to examine if the observed difference in CAVII KO and WT spine architecture is age- and/or area specific, and to see if the absence of CAVII is reflected as increased motility or turnover of the thin, immature-like spines.

So far, a structural role has not been reported for any of the CA_i_s, although studies on CA-related protein VIII KO mice have shown that deletion of this catalytically inactive isoform causes abnormalities in parallel fiber and Purkinje cell synapses (Hirasawa et al., 2007). The combined role of CAVII in ion-regulatory and morphogenic function reported here, bears much resemblance to that described for the K-Cl cotransporter (KCC2) that promotes spine development by a transport-independent interaction with the cytoskeleton (Kaila et al., 2014a). Furthermore, both of these ion-regulatory proteins are involved in the qualitative and quantitative change of GABAergic transmission during brain development (Ruusuvuori et al., 2010; Kaila et al., 2014b). In mature neurons, upon intense GABA_A_ receptor activation, synergistic activity of K-Cl transport and CA activity is able to render GABAergic signalling excitatory and even pro-convulsant (Kaila et al., 1997; Ruusuvuori et al., 2004; Viitanen et al., 2010; Avoli and de Curtis, 2011).

The pH sensitivity of actin polymerization and depolymerization in spines is intriguing. Diering and colleagues (2011) reported an NHE5-dependent increase in spine pH, developing over tens of minutes after chemical long-term potentiation induction in cultured rat hippocampal neurons. Rapid, depolarization-induced acid transients in dendrites have been detected in Purkinje cells (Willoughby and Schwiening, 2002). However, the influence of effects of this kind on spines remains to be explored in more detail. Notably, the effect of pH on actin polymerization and depolymerization is dictated by several factors. Actin self-assembly *in vitro* is enhanced by protons (Wang et al., 1989; Heath et al., 2013) but in the intracellular milieu the pH-sensitivity of actin-binding proteins, such as cofilin and gelsolin, bring additional players to F-actin dynamics (Yonezawa et al., 1985; Azuma et al., 1998; Frantz et al., 2008). Nevertheless, actin-associated, spine-targeted CAVII expression is an exciting observation as synaptic activity is known to evoke long-term structural changes in spine size and morphology (Sala and Segal, 2014), as well as large transient changes in ionic concentrations within the spine (Rose et al., 1999; Brini et al., 2017). The present study shows that CAVII is optimally localized not only to separately modulate, but also to provide a link between F-actin dynamics and activity-dependent pH transients, within spines, thereby identifying a novel mechanism of morphofunctional plasticity.

## Materials and Methods

### Animal experiment ethics

All experiments involving animals were conducted in accordance with the European Directive 2010/63/EU, and were approved by the National Animal Ethics Committee of Finland or the Local Animal Ethics Committee, University of Helsinki.

### Neuronal primary cultures, fibroblast cultures and transfections

Hippocampal neuronal cultures were prepared as described previously (Bertling et al., 2012). Briefly, the hippocampi of embryonal day 17 Wistar rat fetuses of either sex were dissected, the meninges were removed, and the cells were dissociated with 0.05 % papain and mechanical trituration. The cells were plated at a density of 100,000 cells/coverslip (diameter 13 mm), coated with Poly-L-Lysine (0.1 mg/ml; Sigma), in neurobasal medium (Gibco) supplemented with B-27 (Invitrogen), L-glutamine (Invitrogen), and penicillin-streptomycin (Lonza). Transient transfections were performed after 13 days *in vitro* (DIV) using Lipofectamine 2000 (Invitrogen), as described earlier (Hotulainen et al., 2009). Prior to all experiments, we confirmed that cultures formed a dense network of neurons, ensuring the availability of a proper synaptic network. The neurons were imaged after fixation with 4 % paraformaldehyde (PFA). Fibroblasts were maintained in DMEM supplemented with 10 % fetal bovine serum (Hyclone), 2mM L-glutamine (Invitrogen) and penicillin-streptomycin (Lonza). Cells were transfected with Superfect (Qiagen) or Turbofect (Thermo Scientific)-transfection reagent according to manufacturer’s instructions for 24 h and either imaged live or after fixation with 4 % PFA.

### Plasmid constructs

pEGFP-N1 (EGFP) and mCherry-C1 (mCherry) plasmids were purchased from Clontech Laboratories, Inc. Human GFP-β-actin (Dopie et al. E544-E552) and mCherry-β-actin plasmids were gifts from Maria Vartiainen (University of Helsinki, Finland) and Martin Bähler (University of Münster, Germany), respectively. Constructs containing full-length CAII and CAVII (human isoform 1) coding sequences were obtained from ImaGenes (human CAII and human CAVII including start and stop codon; OCAAo5051H1054 and OCAAo5051E0588, respectively) and GeneCopoeia (human CAII without stop codon and mouse CAVII including start and stop). Constructs encoding CAVII-R223E and CAVII-H96/98C were generated using site-directed mutagenesis (Phusion high fidelity PCR, ThermoFisher) and the correct sequence of PCR amplified sequences was confirmed by full-length sequencing of both strands (DNA Sequencing and Genomics Laboratory, Institute of Biotechnology, Helsinki). More complex mutants containing multiple nucleotide exchanges were commercially synthesized (GenScript). All coding sequences were either available as Gateway entry vectors or subcloned into pDONR or pENTR vectors using the Gateway technology (LifeTechnologies). Expression constructs encoding N- or C-terminal fusion proteins of CAII or CAVII and various reporter proteins (EGFP, dsRed, mCherry) were generated using the Gateway technology and appropriate destination vectors. To allow for stable expression in cell cultures and neurons *in vivo*, all destination vectors contained the CMV/chicken beta-actin gene (CAG) promoter (Niwa et al., 1991). The pCAG-EGFP plasmid was a gift from Connie Cepko (Addgene plasmid # 11150) and served as a control (Matsuda and Cepko, 2004).

### Expression level quantification in NIH3T3 cells

NIH3T3 cells were seeded on 24-well plates with Poly-L-Lysine-coated coverslips at a density of 60,000 cells/well, and on 6-well plates at a density of 300,000 cells/well. The following day, the cells were transfected with the CAVII/II-EGFP constructs using Turbofect transfection reagent (Thermo Scientific) according to the manufacturer’s instructions. For immunofluorescence, the cells were fixed 24 h after transfection with 4 % PFA for 30 min, and stained with DAPI (1:10,000 for 2 min) (Fig. S4A). For Western blotting, the cells were washed once with ice-cold phosphate buffered saline (PBS) and collected in 150 µl radioimmunoprecipitation assay (RIPA) buffer supplemented with protease inhibitors (cOmplete, Roche) 24 hours after transfections. 10 µg of the lysates were separated by SDS-PAGE and blots were probed with a 1:2,000 dilution of mouse anti-GFP antibody (Clontech), followed by 1:5000 anti-mouse Starbright Blue 700 (Bio-Rad) (Fig. S4B) and 1:5000 anti-actin-rhodamine (Bio-Rad) as a loading control. The blots were imaged with ChemiDoc MP (Bio-Rad). Quantification was done with ImageJ, and expression levels of the constructs were shown relative to the WT EGFP-CAVII expression level, which was set at one.

### Actin filament staining with phalloidin in fibroblasts

Fibroblasts were permeabilized with 0.1 % tritonX-100 in PBS. Filamentous actin was visualized with Alexa fluor 488- or Alexa fluor 594 -phalloidin (for 30 min, Invitrogen, Molecular probes).

### Actin filament visualization in cultured neurons

Neuronal cultures were co-transfected with a mCherry-actin and EGFP-CAII/CAVII constructs on DIV13 and fixed with 4 % PFA (30 min) 24 h after transfection

### Imaging

Fixed NIH3T3 fibroblasts were imaged under epifluorescence illumination using an upright Axio Imager.M2 microscope equipped with a 63x 1.4NA objective and with an Apotome 2 structured illumination slider (all from Zeiss). Images were acquired with a black and white CMOS camera (Hamamatsu ORCA Flash 4.0 V2) and ZEN 2 software (Zeiss). For quantification of co-localization of the EGFP-tagged proteins with 594-Phalloidin in fibroblasts we analyzed the pixel intensities along a virtual line across the cell, excluding the nucleus (ImageJ, http://imagej.nih.gov/ij). The placing of the line was done using the “actin channel” and the experimenter was blinded to the transfections. We plotted the pixel intensities of both channels against each other and calculated the Pearson’s correlation coefficient (GraphPad Prism 7). The dendritic spines of cultured hippocampal neurons were imaged using a Zeiss LSM 710 upright confocal microscope (63x 1.3NA objective) or an Axio Imager.M2 microscope (63x 1.4NA objective, Apotome 2 structured illumination slider). Image files were processed with ZEN 2012 (Carl Zeiss Microscopy GmbH), ImageJ 1.46r and Photoshop CS4 (Adobe).

### SEC-MALLS

The size-exclusion chromatography-coupled multi-angle static laser light scattering (SEC–MALLS) was used for characterisation of the oligomerization and monodispersity of CAVII essentially as described in (Karki et al., 2018). The measurements were done at 0.5 ml/min over an S-200 Superdex 10/300 column (GE Healtcare) in 1x PBS with a HPLC system (Shimadzu) and a MiniDAWN TREOS light scattering detector, and Optilab rEX refractive index detector (Wyatt Technology Corp.). Data were then analysed with ASTRA 6 software (Wyatt Technology Corp.). CAVIII protein was analysed at 33 µM concentration in 50–100 μl volume.

### Spine analysis

For analysis of spine density and morphology in fixed rat cell cultures, serial image files corresponding to *z*-stacks of 20–30 optical sections per dendritic segment were taken. Only healthy looking cells (no beading of dendrites or other signs of decreased wealth) with spines were imaged and included to analysis. NeuronStudio, a software package specifically designed for spine detection and analysis (Rodriguez et al., 2008) was used to analyze spine density. The detailed analysis of spine classes was performed as described in (Bertling et al., 2012). Classification of spines was done by using rules defined by (Rodriguez et al., 2008) and verified manually. For the plot, spines were divided to three groups: spines with head (Neurostudio: mushroom and stubby), filopodia/thin spines (Neurostudio: thin) and spines with abnormal morphology (Neurostudio: other), latter including all spines with branches, long thick protrusions or otherwise morphology not classified in any common spine classes. The mean value (+ SEM) of separate images is shown.

### Latrunculin B assay

For the Latrunculin B assay fibroblasts were transfected with dsRed or dsRed-CAVII 24 h prior to treatment with 5 µM Latrunculin B in DMSO. Cells were fixed after 0, 2, 5, 10, and 30 minutes of Latrunculin B treatment and stained with phalloidin-488. Control cells were treated with DMSO for 60 min and stained with phalloidin-488. We made three independent replicates of such experiments. A hundred transfected cells from each time point from each experiment were categorized either as “normal”, “some shape/F-actin left” or “round” (example cells for the three categories are depicted in Figure 3A,B, upper panels).

### CAVII biochemistry: pull down assay, measure of enzymatic activity, bundling (In vitro TIRF) assay

Actin co-sedimentation assay was carried out in 20 mM Hepes pH 7.4/6.5, in the presence of 0.2 mM DTT. Mouse CAVII (produced as a secreted protein with a C-terminal His-tag in the CHOEBNALT85 cell line) was stored in PBS but the buffer was changed to Hepes (pH 7.4/6.5) before the experiment. Lyophilized powder of CAII (Sigma) was reconstituted in MilliQ and diluted in Hepes-buffer pH 7.4/6.5 to 33 μM. ZnCl_2_ (1 μM) was added to CAVII/CAII one hour before incubation with actin. β/γ-G-actin (0, 1, 5, 10 and 15 μM) was pre-polymerized in Hepes-buffer pH 7.4/6.5 by addition of 1/10 of 10x-initiation mixture (1 M KCl, 10 mM EGTA, 50 mM MgCl_2_, 2.5 mM ATP and 20 mM Hepes pH 7.4/6.5) for 30 min at room temperature. CAVII or CAII (1 μM) was added to polymerized actin, gently mixed and incubated for another 30 min at room temperature. Actin filaments were sedimented by centrifugation for 30 minutes at 20S°C in a Beckman Optima MAX Ultracentrifuge at 353,160 × g in a TLA100 rotor. Equal proportions of supernatants and pellets were run on 13.5 % SDS-polyacrylamide gels, which were stained with Coomassie Blue. The intensities of β/γ-actin and CAVII/CAII bands were quantified with QuantityOne program (Bio-Rad), analyzed and plotted as CAVII/CAII bound to actin (μM, CAVII/CAII in pellet) against actin. The mCAVII-actin co-sedimentation assay was repeated three times for each pH value and averaged curves were presented (± SEM).

*In vitro* TIRF imaging was performed as previously described (Suarez et al., 2011) but the muscle actin was substituted with non-muscle actin (Cytoskeleton), prepared according to the manufacturer’s instructions, and non-muscle Rhodamine actin (Cytoskeleton) was used for labeling the filaments. A mixture of 0.5 µM unlabeled and 0.05 µM Rhodamine labelled non-muscle actin was polymerized in the presence of 1.12 µM mCAVII or with an identical volume of PBS as a control in the nominal absence of CO_2_/HCO_3_^-^ (three independent repeats for both treatments). Images were captured with Nikon Eclipse Ti-E N-STORM microscope, equipped with Andor iXon+ 885 EMCCD camera and 100x Apo TIRF oil objective (NA 1.49), a 150 mW 561 nm laser line was used for visualization of Rhodamine actin. Actin filament polymerization was followed (images every 10 sec) until the imaging field was full with filaments (typically around 30 - 40 min). Bundling was quantified by measuring the mean relative fluorescence intensity of a cross-section for an individual filament (*n* = 10 - 11 filaments per repeat, ImageJ) and actin fiber length (*n* = 3 – 4 per repeat) at three time points (0, 5, and 23 minutes). The data was analyzed using a general mixed model with time as a within unit factor and the presence of CAVII as a between unit factor, experiment repeats were included as a covariate.

### In utero electroporation and slice preparation

The following modifications to the rat IUE protocol described in (Fiumelli et al., 2013) were applied for mice. Timed-pregnant ICR mice with E14.5 embryos were given Temgesic (0.05-0.1 mg/kg, s.c.) and anesthetized with isoflurane (4.2 % induction, 2.5 % during surgery). All embryos were injected with 1.25 µl plasmid DNA solution (3-4 µg/µl EGFP-CAVII or EGFP-CAII construct in 0.9 % NaCl and 0.1 % Fast Green). Electroporation was done with 5 mm diameter circular electrodes (Sonidel Limited) with five pulses (40 - 45 V, 50 ms duration at 100 ms intervals), delivered with a square-wave generator (CUY21vivo SC, Sonidel Limited). Detection of EGFP-CAII and EGFP-CAVII was done on 50 µm coronal cryosections from fixed brains (P40 mice were transcardially perfused under terminal anesthesia with 4 % PFA, over-night postfixation in 4 % PFA) with a Zeiss LSM 710 upright confocal microscope.

### Cortical layer 2/3 spine analysis and mEPSC recordings

#### Post Hoc iontophoretic injection of Lucifer Yellow

Male WT and CAVII KO mice were terminally anesthetized at P34 - P37 by an intraperitoneal injection of pentobarbital (100 mg/kg) and perfused transcardially first with saline, followed by 4 % PFA and 0.125 % glutaraldehyde solution (pH 7.4). Brains were removed and postfixed for 2 h in 4 % PFA. Coronal sections of 200 µm thickness were cut on a vibratome in ice-cold PBS (pH 7.4). Coronal sections were pre-stained for 10 min with methylene blue, which allows the visualization of neuronal somata, mounted into an injection chamber, and placed on the fixed stage of a Zeiss microscope equipped with a micromanipulator. Layer 2/3 pyramidal neurons were loaded iontophoretically with a 0.4 % Lucifer yellow solution (Sigma-Aldrich, St. Louis, MO) using sharp micropipettes and a negative current of 70 nA until the dendrites were fluorescing brightly. For each animal, neurons were labelled from 2-3 slices.

### Immunohistochemistry

The Lucifer Yellow injected slices were preincubated for 1 h in a PBS solution containing sucrose (5 %), bovine serum albumin (2 %), Triton X-100 (1 %) and sodium azide (0.1 %), followed by 48 h at room temperature with the anti-LY antibody (rabbit IgG, Cat.No. A5750, Invitrogen, Carlsbad, CA; 1:4,000 dilution). Slices were then rinsed in PBS solution and incubated for an additional 24 h with Alexa conjugated secondary antibodies (Invitrogen; 1:1,000). After mounting the slices were coverslipped using Immumount (Thermo Scientific, Pittsburgh, PA), and stored at +4 °C until analysis.

### Confocal Laser Scanning Microscopy and Image Analysis

Second order dendrites were imaged for spine analysis using LSM700 confocal microscope and 63× oil-immersion objective. Spine analysis was performed on acquired stacks of images using a homemade plug-in written for OsiriX software (Pixmeo, Geneva, Switzerland). This plug-in allows precise spine quantification, individual tagging, and measurement in 3D by scrolling through the z-axis. We defined spines as structures emerging from the dendrites that were longer than 0.4 µm and for which we could distinguish an enlargement at the tip (spine head). Spines head diameters were measured at their largest width in xy-axis on the z-image corresponding to the central axis of the spine head. The difference in spine head width distribution between WT and CAVII KO mice was analyzed using a Wilcoxon rank sum test with continuity correction. Note that for illustration purposes, images presented in figures are maximum intensity projections of z stacks with volume rendering, further treated with a Gaussian blur filter.

### mEPSC recordings and analysis

Male CAVII KO and WT mouse (P30 - P40) were anesthetized with halothane and decapitated. Acute coronal brain slices (400 µm) were cut using Campden vibratome (Campden Instruments 7000 SMZ-2) in ice cold (<4°C) cutting solution containing (in mM) 87 NaCl, 2.5 KCl, 0.5 CaCl_2_, 25 NaHCO_3_, 1.25 NaH_2_PO_4_, 7 MgCl_2_ and 50 sucrose, equilibrated with 95 % O_2_ and 95 % CO_2_ to pH 7.4. Before starting experiments the slices were let to recover for one hour at +34°C in standard solution containing (mM): 124 NaCl, 3 KCl, 2 CaCl_2_, 25 NaHCO_3_, 1.1 NaH2PO_4_, 1.3 MgSO_4_ and 10 D-glucose, (300 ± 5 mOsm). Whole-cell voltage-clamp recordings from layer 2/3 somatosensory cortex pyramidal neurons were obtained with a HEKA EPC-10 amplifier with 20 kHz sampling interval and 4 kHz low-pass filter. Slices were perfused with standard solution (see above, perfusion 3.5 ml/min) and all measurements were done in the presence of 100 µM picrotoxin and 0.5 µM TTX. Temperature in the recording chamber was 32 ± 1°C. The cells were clamped to −65 mV. Borosilicate patch pipette resistances ranged from 3 - 5.5 MΩ when filled with a pipette solution containing (mM) 140 CsMs, 2 MgCl_2_, 10 HEPES liquid junction potential of 13 mV was taken into account). Only cells with a resting membrane potential below −55 mV and stable holding current were included in the analysis. Series resistance was compensated and recordings with unstable series resistance (change > 30 %) were excluded from the analysis. The person who did and analyzed the mEPSC experiments was blind to the genotype. Events were manually detected with minianalysis software (synaptosoft) after 1000 Hz low-pass filtering and with threshold set to 4x RMS noise.

## Acknowledgements

We wish to acknowledge Alexander Alafuzoff for his indispensable help with statistical analysis; Merle Kampura for her expertise in performing *in utero* electroporation and handling cell cultures; Mikko Liljeström and the Biomedicum Imaging Unit (BIU), Helsinki, for kind help and technical assistance with the *in vitro* TIRF imaging experiments. The use of the INSTRUCT-HiLIFE protein crystallization core facility, University of Helsinki, member of Biocenter Finland and Instruct-FI is gratefully acknowledged for assistance in the SEC-MALLS experiments.

## Author contributions

PB, KK, PH and ER conceptualized the study. PB, MB and VP designed constructs. EB, PB, PS, MS, VP, LV, KK, PH and ER designed the experiments. EB, PB, PS, EK, GG, MV, MS, IS and VP collected data. EB, PS, EK, GG, MV, MS, IS, VP and LV analyzed data. PB, KK, PS, VP and PH reviewed and edited the paper. ER wrote the paper.

## Figure legends

**Figure 1 - figure supplement 1.**
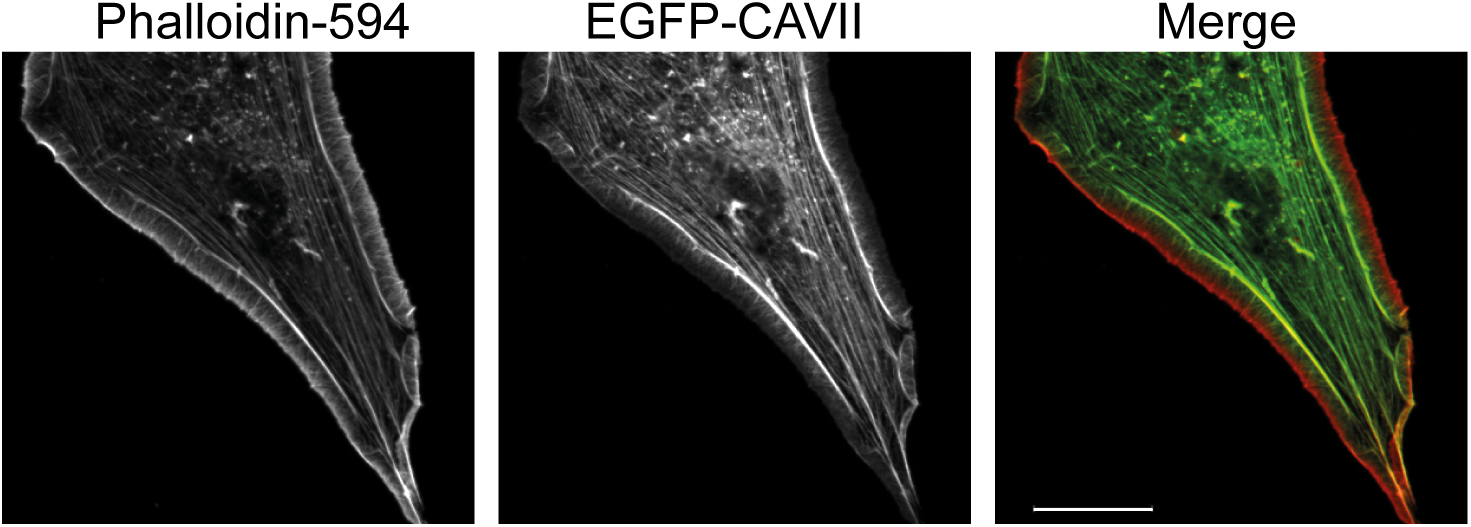
CAVII co-localizes with selective actin filaments. EGFP-CAVII co-localizes with subcellular F-actin structures (visualized with Phalloidin-594) except at the very edges of fibroblast lamellipodia. Scale bar 20 µm.

**Figure 2 –figure supplement 1.**
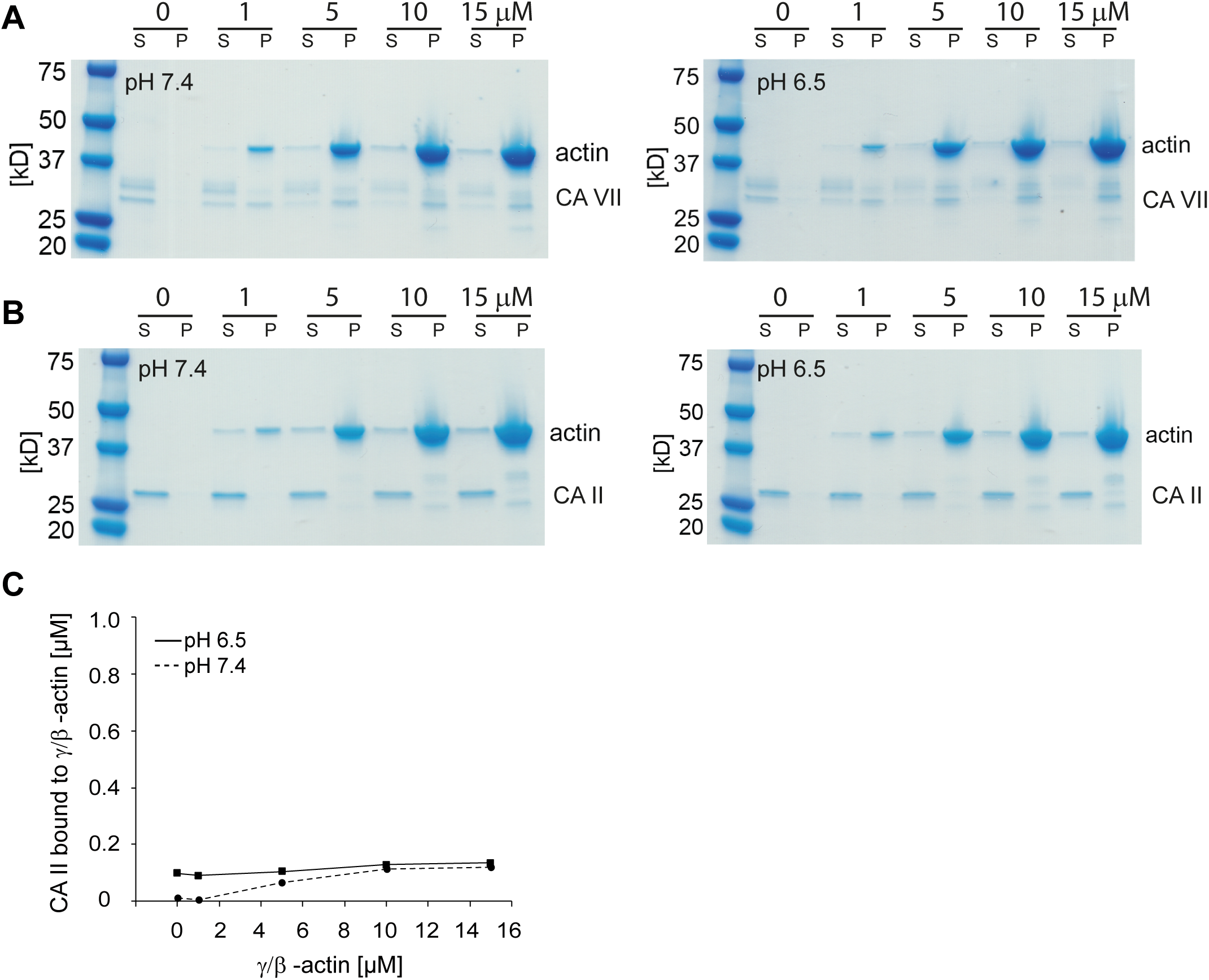
CAVII, but not CAII, co-localizes and directly interacts with F-actin. Actin co-sedimentation assay was carried out at five different concentrations of β/γ-actin and with 1 µM (A) CAVII or (B) CAII at two different pH (7.4 or 6.5). After centrifugation, the supernatant (S) and pellet (P) fractions were separated and resolved by SDS-PAGE. Staining the gels with Coomassie Blue showed that CAVII co-sedimented in the pellets with actin, whereas CAII was found only in the supernatant fraction. CAVII; three repetitions and CAII; one repetition at each of the four actin concentrations/pH. (C) Analysis of the CAII gels confirmed that the isoform does not interact with actin at either pH tested.

**Figure 2 –figure supplement 2.**
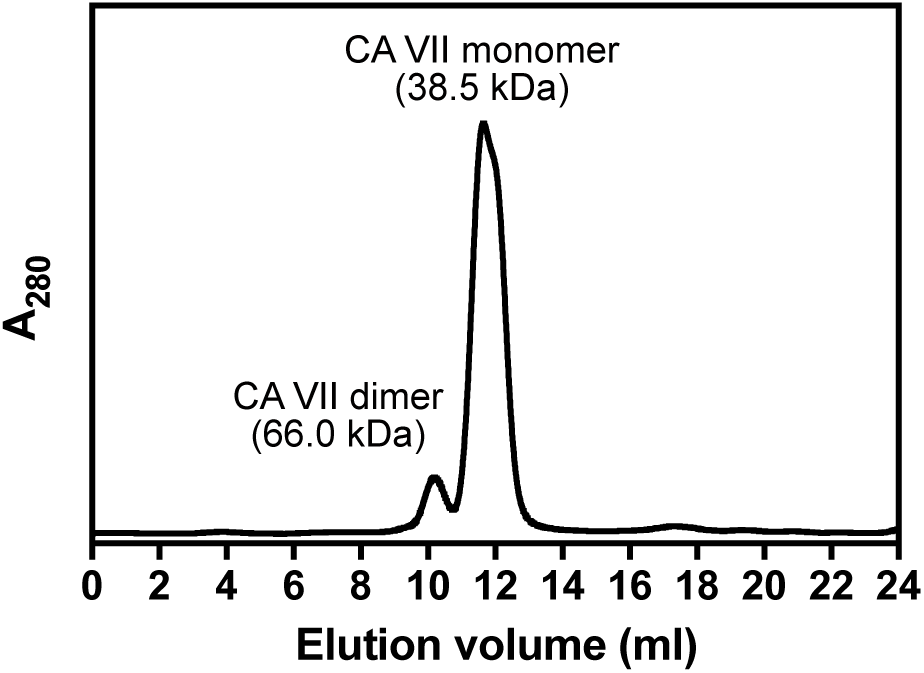
Purified recombinant CAVII exists as a mixture of monomers and dimers. CAVII analytical gel filtration analysis. CAVII was run on Superdex 200 10/300 gel filtration column in TBS at 0.5 ml/min, at protein concentration of 33 µM. The experiment was carried out once and SEC-MALLS data were analyzed using with ASTRA 6 software (Wyatt Technology Corp.) as described in Folta-Stogniew and Williams (1999). A major peak at approximately 12 ml volume eluted and based on multi-angle light scattering had molecular weight of *ca.* 38.5 kDa, matching relatively well with theoretical molecular weight of the monomer (30.5). The molecular weight determined for the minor peak (66,6 kDa) eluting at approximately 10 ml corresponds roughly to a CAVII dimer.

**Figure 2 - Source Data 1** The file shows the values for CAVII and CAII bound to actin (μM, CAVII/CAII in pellet) against actin. These data were used for the quantitative analyses shown in Figure 2A and Figure 2 – figure supplement 1C. The intensity values of β/γ-actin and CAII/CAVII bands were quantified with QuantityOne program (Bio-Rad). All experiments were included in the analysis.

**Figure 2 - Source Data 2** The measured actin fiber lengths (*n* = 3 – 4 filaments per repeat) and the mean relative fluorescence intensity values of cross-sections for individual filaments (*n* = 10 - 11 filaments per repeat, ImageJ) that were used to quantify bundling presented in Figure 2C. The analyzed filaments were chosen randomly and all experiments were included in the analysis.

**Figure 3 - Source Data 1** Source files for WT and CAVII over expressing cells. Excel file contains the counted cells, divided to different categories based the F-actin staining. Averages from the three experiments is shown in Figure 3. Inclusion criteria: all cells expressing dsRed (control) or dsRed-CAVII. We did not exclude any outliers.

**Figure 4 – figure supplement 1.**
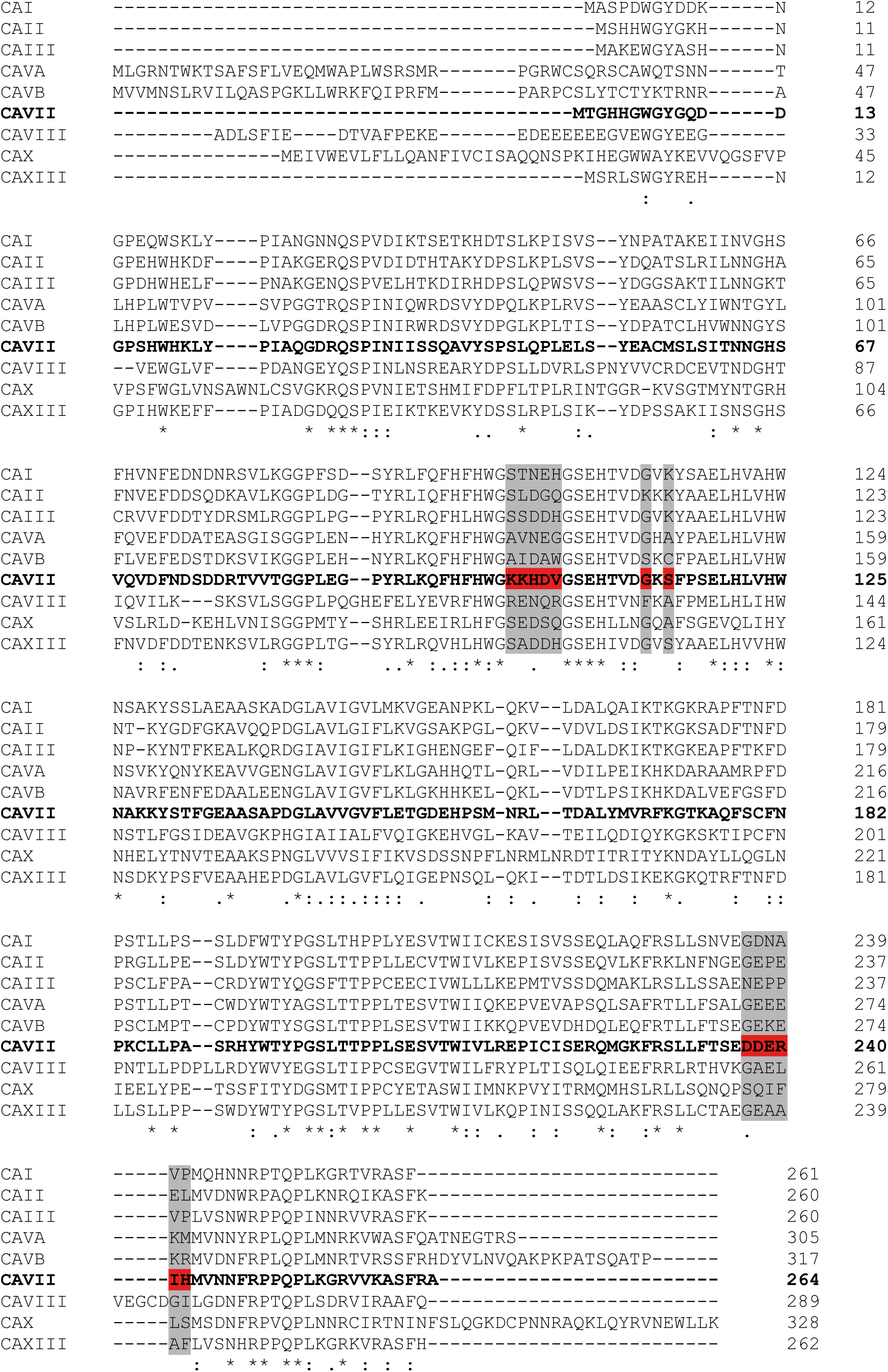
The identified actin interaction sites are unique to CAVII among human cytosolic CA’s. Sequence alignment of the catalytically active (CA I, CAII, CAIII, CA VA, CA VB, CAVII, and CA XIII) and catalytically inactive (CAVIII and CA X) human cytosolic CA protein sequences generated using the Clustal O (1.2.1) multiple sequence alignment. The amino acids that were characterized as part of a putative actin binding site of CAVII (highlighted in red) are not conserved in the other cytosolic CA isoforms (highlighted in gray). An asterisk below the aligned sequences indicates fully conserved residues, a colon indicates residues with strongly similar properties (scoring > 0.5 in the Gonnet PAM 250 matrix) and a period indicates residues with weakly similar properties (scoring ≤ 0.5 in the Gonnet PAM 250 matrix).

**Figure 5 – figure supplement 1.**
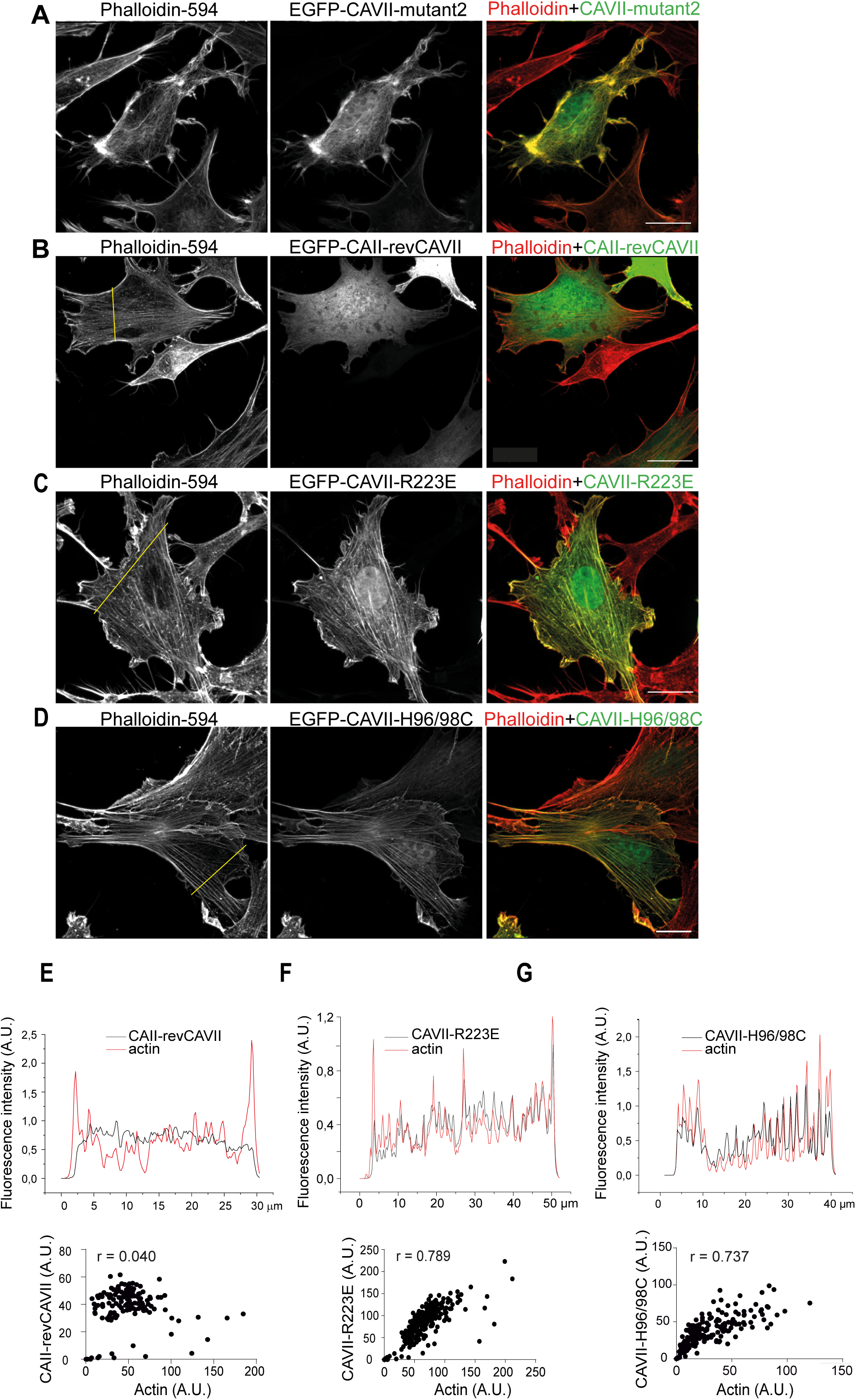
Subcellular localization of the chimeric CA fusion proteins EGFP-CAVII-mutant2, EGFP-CAII-revCAVII, EGFP-CAVII-R223E and EGFP-CAVII-H96/98C in fibroblasts. (A)Transfection with EGFP-CAVII-mutant2 modified cellular F-actin structures in a similar manner than CAVII. (B) Introduction of KKHDV and DDERIH motifs to CAII (EGFP-CAII-revCAVII) did not affect the diffuse cytosolic localization of the isoform II. (C) EGFP-CAVII-R223E and (D) the catalytical loss-of-function mutant EGFP-CAVII-H96/98C co-localized with F-actin. F-actin is visualized with Phalloidin-594 in A - D. The normalized fluorescence emission intensity profiles of (E) EGFP-CAII-revCAVII (F) EGFP-CAVII-R223E, and (G) EGFP-CAVII-H96/98C (black lines) and F-actin (red line). The yellow line in B – D indicates the cross-section of the cell from which the pixel intensities were measured. Analysis of co-localization is shown in lower panels of E - G. Representative single-cell pixel intensities of EGFP and phalloidin-594 channels were plotted against each other and the Pearson’s correlation coefficient value (r) was calculated. *n* = 3-4 independent transfections/construct. Scale bars in A – D is 20 µm.

**Figure 5 – figure supplement 2.**
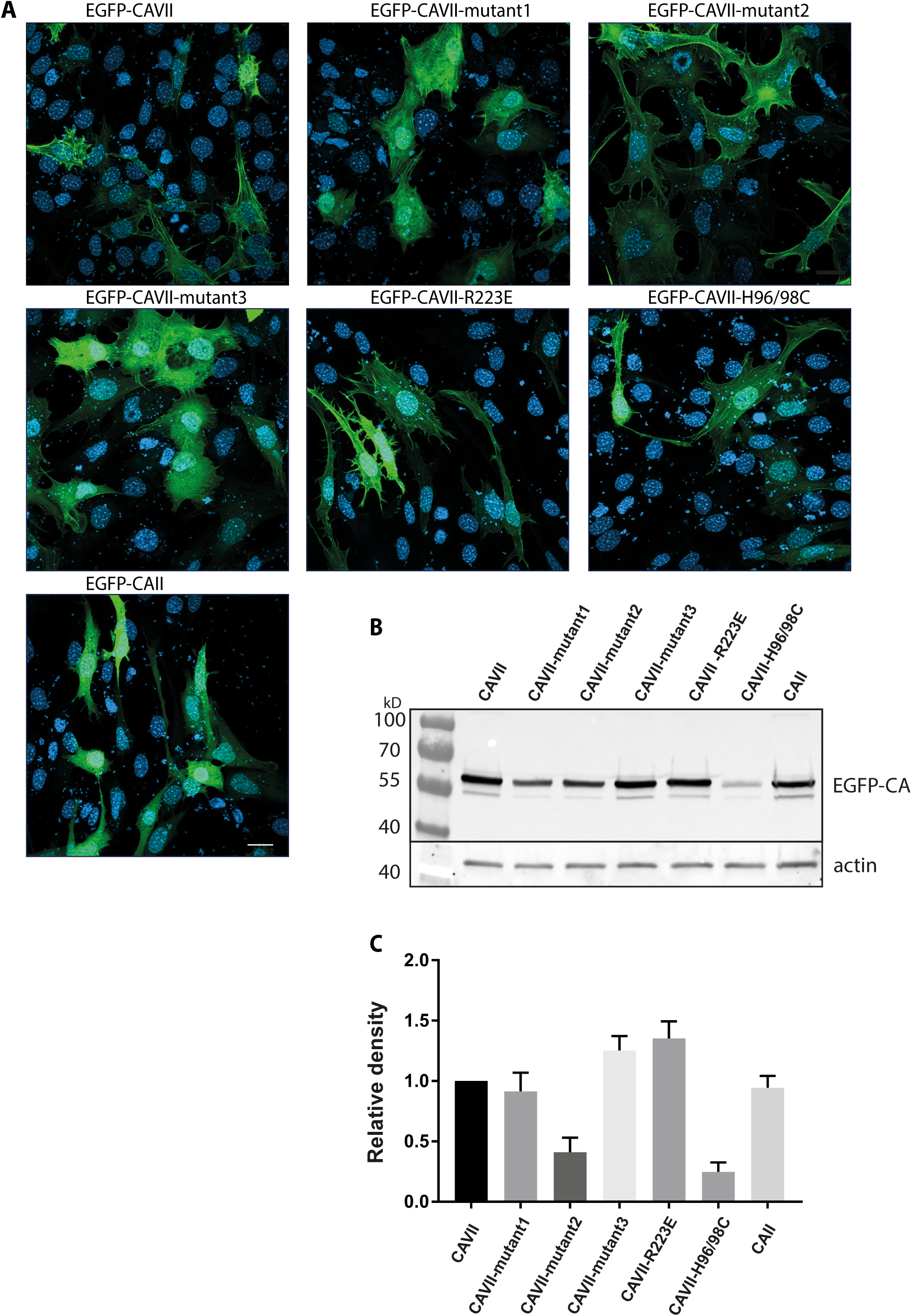
Expression levels of the different EGFP fusion proteins in fibroblasts. (A) Representative images of NIH3T3 cells transfected with the EGFP-tagged CA constructs. Cells were fixed 24 hours after transfection and nuclei were stained with DAPI. *n* = 2 - 3 independent transfections for each construct. Scale bar 20 µm. (B) A representative Western blot showing the expression levels of EGFP-CA fusion proteins in 10 µg of lysate collected 24 hours after transfection. The EGFP-tagged CA fusion proteins are visible at approximately 60 kDa, and the β-actin loading control is visible at 42 kDa. (C) Quantification of the fusion proteins expression levels in NIH3T3 cells. Expression level of EGFP-CAVII was set at 1 for each Western blot. *n* = 5 independent transfections for all fusion proteins except for CAVII-mutant2, for which *n* = 4. Transfections efficacy of different fusion proteins did not differ significantly from each other (one-way ANOVA with Dunnett’s multiple comparisons test).

**Figure 5 – Source Data 1/** This spreadsheet contains Pearson’s correlation coefficients calculated for EGFP-tagged CA constructs vs. F-actin for all analyzed NIH3T3 fibroblasts in Figure 5D and E. Each individual experiment consists of 1-4 transfected wells. The person who analyzed the co-localization and calculated Pearson’s coefficient values was blind to the transfection.

**Figure 5 – Source Data 2/** The quantified expression levels of the different WT and mutant CA’s in NIH3T3 fibroblasts. Averages from the five experiments is shown in Figure 5-figure supplement 2. The CA-EGFP band area was quantified (using Image J) as a percentage of the CAVII WT-EGFP band, which was set at 1 on each blot. The actin band area was quantified as a percentage of the actin band on the CAVII WT-EGFP lane, which was set at 1 on each blot. Finally, the relative density for each band was calculated by dividing the band density for CA-EGFP by the band area for actin on the same lane. One lane (mutant2 on 18.1.2019) was excluded because of a mistake in the cell transfection: the actin band was normal but there was no EGFP band on the lane.

**Figure 6 – figure supplement 1.**
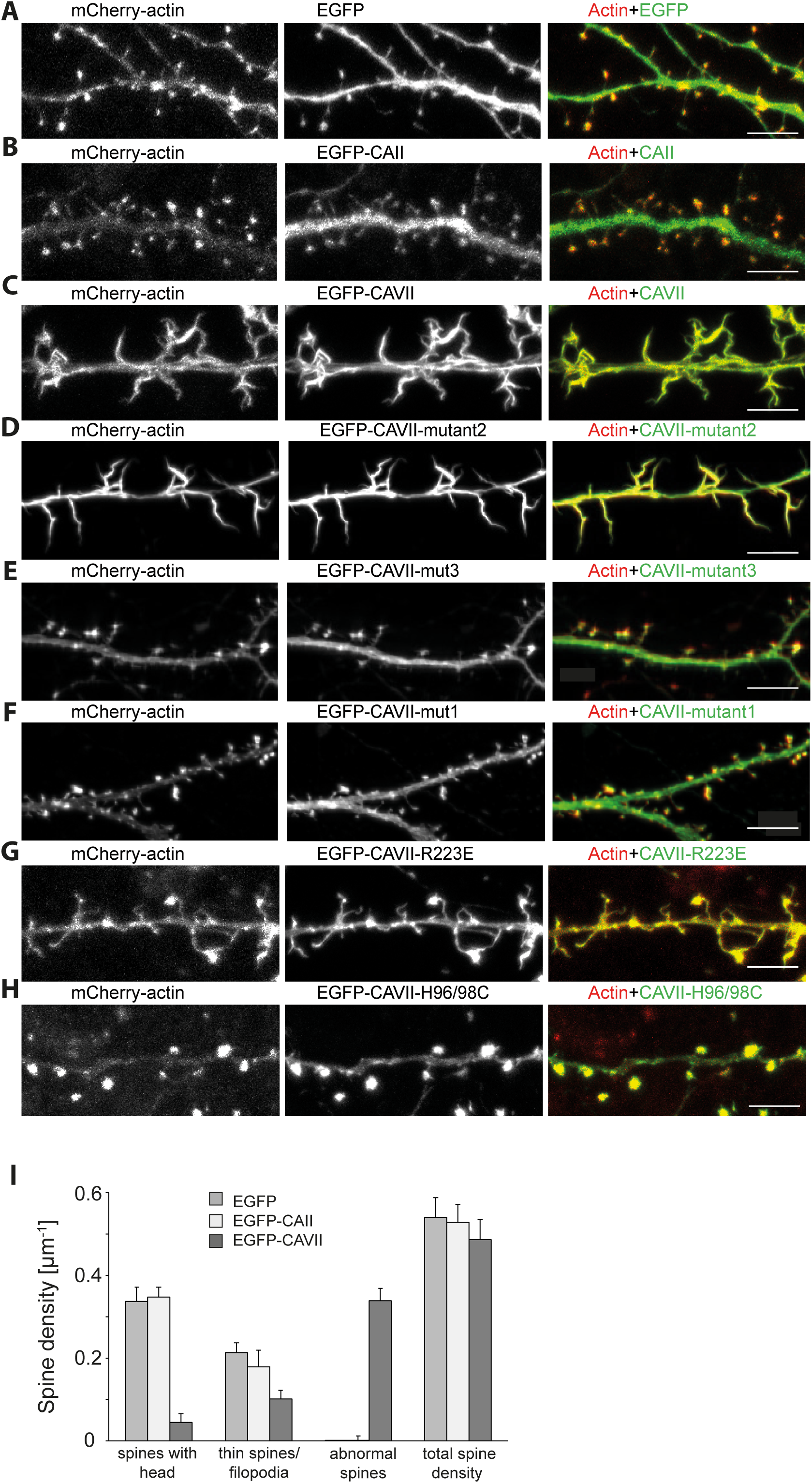
Localization of WT and chimeric CA EGFP-fusion proteins along the dendritic shaft and in spines in cultured hippocampal neurons. (A) Control experiments with neurons co-expressing mCherry-actin and EGFP. (B) Neuron transfected with mCherry-actin and EGFP-CAII. Compared to the spine-targeted mCherry-actin, CAII localizes more diffusely along dendritic shafts and spines. Both (C) EGFP-CAVII and (D) EGFP-CAVII-mutant2 show a highly overlapping localization with mCherry-actin and disruption of dendritic spine morphology. Spines were replaced by thick, filopodia-like dendritic protrusions, which lack spine heads. The loss-of-function constructs (E) EGFP-CAVII-mutant3 and (F) EGFP-CAVII-mutant1 are more homogenously present in both dendrites and spines. (G) EGFP-CAVII-R223E and (H) the catalytically inactive EGFP-CAVII-R223 98C showed overlapping localization with mCherry-actin. Scale bar 5 µm (A – C, G – H), 10 µm (D – F).

**Figure 6 – figure supplement 2. EGFP-CAVII expression disrupted normal spine morphology in cultured neurons** EGFP-CAVII expression disrupted normal spine morphology in cultured neurons: control, only mCherry-actin: spines with head 0.34±0.04, thin spines/filopodia 0.21 ± 0.02, abnormal spines 0.00 ± 0.00, total 0,54±0.05 spines/µm; *n* = 10 cells, 509 spines, 973 µm analyzed dendrite; EGFP-CAII: spines with head 0,35 ± 0.04, thin spines/filopodia 0.17 ± 0.02, abnormal spines 0.00 ± 0.00, total 0.53 ± 0.06 spines/µm; *n* = 10 cells, 535 spines, 992 µm analyzed dendrite; EGFP-CAVII: spines with head 0.05 ± 0.02, thin spines /filopodia 0.10 ± 0.02, abnormal spines 0.34 ± 0.03, total 0.49 ± 0.05 spines/µm; *n* = 10 cells, 493 spines, 10256 µm dendrite. Analyzed cells were pooled from two independent experiments. Data are presented as mean + SEM.

**Figure 6-Source Data 1** Source files for spine analysis of cultured neurons expressing only mCherry-actin, or mCherry-actin with EGFP-CAII or EGFP-CAVII. Excel file contains the calculated spine densities for each analyzed cell used for the quantitative analyses shown in Figure 6–figure supplement 2. Inclusion criteria: all healthy, pyramidal neuron looking cells expressing moderate amount of mCherry and co-expressed EGFP-construct. We did not exclude any outliers.

**Figure 7-Source Data 1/Source files for WT and CAVII KO spine analysis** These Excel file contains the calculated spine densities and spine head size distribution used for the quantitative analyses shown in Figure 7 B and C. Inclusion criteria: Layer 2/3 pyramidal neurons in the somatosensory cortex with Lucifer Yellow (LY) -filled dendritic tree. Exclusion criteria: LY-filled neurons outside layer 2/3 and neurons where LY-injections failed (leading to partially filled dendritic arbor). We did not exclude any outliers. All data were counted and the person who analyzed spine density/spine head size was blind to the genotype.

## Rich Media Files

**Figure 2, Videos 1 and 2. Actin polymerization, visualized with Rhodamine, in the absence (video 1) and presence of CAVII (video 2).** Time-lapse images were taken every 10 s. The total duration of both videos is 23 min and they are displayed at a rate of seven frames/second.

